# Defining the *Xanthomonas euvesicatoria* type II-secreted effector arsenal reveals key roles in bacterial nutrition and diversity across species

**DOI:** 10.64898/2026.06.19.733414

**Authors:** Samuel Goll, Theresa Staps, Kristina S. Munzert-Eberlein, Lilly Hafen, Akash K. Shivhare, Iliyana Kraleva, Susanne Matschi, Stephanie Krüger, Timo Engelsdorf, Daniela Büttner, Jessica L. Erickson

## Abstract

- The type II secretion (T2S) system is conserved across the *Xanthomonas* lineage, yet its contributions to pathogenicity and secreted protein repertoires are poorly defined. We demonstrate that T2S systems in *Xanthomonas* pathovars with divergent hosts and lifestyles are required for disease.
- Quantification of the plant cell wall composition during infection by *Xanthomonas euvesicatoria* (*Xe*) revealed a T2S-dependent depletion of galacturonic acid, suggesting T2 effector (T2E)-mediated cell wall remodeling.
- Using an *in planta* label-free proteomics approach, we identified two known and 20 new *Xe* T2Es from tomato apoplast, most annotated as cell wall-degrading enzymes and proteases. *In vitro* growth assays showed a T2S-mediated utilization of plant cell wall polysaccharides and proteins by *Xe*, as well as by *Xanthomonas axonopodis* pv. *glycines* (*Xag*) and *Xanthomonas campestris* pv. *campestris* (*Xcc*). Intriguingly, while families of cell wall-degrading enzymes are relatively well conserved across these three strains, protease repertoires have diversified significantly, suggesting more strain-specific functionalities.
- Our methodology establishes a framework for T2E discovery, enabling future functional dissection of this understudied effector class and its crosstalk with other bacterial virulence factors.

## Introduction

Plant-pathogenic bacteria infect a wide range of economically important crop plants and thus cause severe yield losses worldwide (Savary *et al*., 2019). Bacterial infection and colonization of plants often depend on a variety of virulence factors, which are secreted into the extracellular milieu or transported into host cells. In Gram-negative bacteria, delivery of virulence factors depends on specialized protein secretion systems, classified into at least nine types (Costa *et al*., 2015; Denise *et al*., 2020; Jamali *et al*., 2025). Among these, the type III secretion (T3S) system is often essential for pathogenicity and translocates type III effector proteins (T3Es) directly into eukaryotic cells (Wagner *et al*., 2018). T3Es reprogram various host processes including signalling pathways, protein turnover or gene expression to promote bacterial survival and multiplication (Büttner, 2016; Zhang *et al*., 2022; Cai *et al*., 2023). Many T3Es suppress plant defense responses, which are activated upon recognition of conserved pathogen-associated molecular patterns (PAMPs) by corresponding pattern recognition receptors (PRRs) (DeFalco & Zipfel, 2021; Ngou *et al*., 2022; Ramirez-Zavaleta *et al*., 2022). This pattern-triggered immunity (PTI) provides a basal resistance against non-specialized pathogens (Jones & Dangl, 2006). As a second layer of plant immunity, T3Es are recognized by intracellular NLR (nucleotide-binding leucine-rich repeats) proteins in resistant plants, triggering effector-triggered immunity (ETI) and often local programmed cell death (Jones & Dangl, 2006; Nivaskumar & Francetic; Ngou *et al*., 2022).

While T3Es promote bacterial colonization from inside the plant cell, successful infection also requires that the pathogen modifies the plant apoplast - the intercellular space between cell membranes including the xylem (Darino *et al*., 2022). A major physical obstacle to bacterial infection in the apoplast is the plant cell wall, a structurally complex barrier which also represents a potential source of carbon and nutrients upon degradation (Fatima & Senthil-Kumar, 2015; Munzert & Engelsdorf, 2025; Pinto *et al*., 2025). The primary cell wall consists of cellulose microfibrils cross-linked by hemicelluloses such as xylan and xyloglucan and embedded in a pectin matrix (Cosgrove, 2024). Cellulose is composed of β-1,4-linked glucose chains and forms the load-bearing framework, while pectin, which is rich in galacturonic acid residues, regulates wall porosity and extensibility (Scheller & Ulvskov, 2010; Obomighie *et al*., 2025). Plant cell walls also contain a substantial amount of protein (Tan *et al*., 2013). This structural complexity hampers the direct utilization of plant cell wall components by pathogens. Given that the plant apoplast is a nutrient-poor environment, many pathogens rely on secreted proteins including cell wall-degrading enzymes (CWDEs) for nutrient acquisition. CWDEs are often secreted by the type II secretion (T2S) system, which is present in many Gram-negative plant- and animal-pathogenic bacteria (Korotkov & Sandkvist, 2019; Naskar *et al*., 2021). T2S systems operate via a two-step mechanism: protein substrates first cross the inner membrane via the general secretory (Sec) or twin-arginine translocation (Tat) pathways, fold in the periplasm, and are then transported across the outer membrane (OM) through the secretin channel (Costa et al., 2015; Korotkov & Sandkvist, 2019). T2S systems contribute to virulence of strains belonging to some of the most damaging plant-infecting bacterial genera, including *Pseudomonas*, *Ralstonia*, *Dickeya*, *Pectobacterium*, *Erwinia*, *Xylella*, and *Xanthomonas* (Yaowei Kang, 1993; Ray *et al*., 2000; Toth *et al*., 2003; Tsujimoto *et al*., 2008; Szczesny *et al*., 2010; Condemine & Le Derout, 2022; Ingel *et al*., 2023). Some necrotrophic pathogens such as *Dickeya* and *Pectobacterium*, secrete pectate lyases to degrade the plant cell wall, thus facilitating both nutrient acquisition and tissue maceration (Fatima & Senthil-Kumar, 2015). In contrast, a direct contribution of type II-secreted proteins, hereafter referred to as type II effectors (T2Es), to nutrient acquisition during apoplast colonization has not yet been demonstrated for hemibiotrophic pathogens.

In addition to potential roles in nutrient acquisition, T2Es may also modulate plant immunity, as was described for extracellular proteins from animal-pathogenic bacteria (Ciaston *et al*., 2022). Notably, a recent study showed that a type II-secreted subtilase from commensal *rhizobacterium* can interfere with plant defense by cleaving the PAMP flg22, which is a derivative of bacterial flagellin and elicits PTI (Eastman *et al*., 2024). Conversely, T2Es themselves can act as PAMPs, thereby triggering rather than suppressing plant defense responses (Dora *et al*., 2022). In addition, cell wall-derived fragments resulting from T2E actions can also serve as damage-associated molecular patterns (DAMPs) and activate PTI responses upon recognition by the plant immune system (Vorholter *et al*., 2012; Molina *et al*., 2024). Cell wall degradation is thus a double-edged sword that simultaneously supports potential nutrient acquisition and risks triggering host defenses. These diverse and sometimes contrasting activities highlight the functional complexity of T2Es and their broad relevance to bacterial colonization of the apoplast. Nevertheless, the specific contributions of individual T2Es to bacterial survival and the full extent of their functional repertoire remain largely unresolved.

*Xanthomonas* provides a compelling model to address these questions, as it is a genus that infects more than 400 economically important plant species worldwide and causes significant agricultural losses (An *et al*., 2020; Timilsina *et al*., 2020). Notably, while some xanthomonads do not possess a T3S system, the T2S system is conserved in all sequenced strains (Alvarez-Martinez *et al*., 2021), indicative of its fundamental importance for members of this genus. *In vitro* studies demonstrate that *Xanthomonas* can degrade cell wall polysaccharides (Szczesny *et al*., 2010; Vorholter *et al*., 2012; Solé *et al*., 2015) and metabolize xylan-derived sugars as a carbon source (Dejean *et al*., 2013; Vieira *et al*., 2021). Cell wall sugar mobilization can also fuel *Xanthomonas* proliferation in citrus leaves (Phan *et al*., 2025).

In the model strain *Xanthomonas euvesicatoria* (*Xe*), the causal agent of bacterial spot disease in pepper and tomato (An *et al*., 2020; Timilsina *et al*., 2020), the T2S system is required for full virulence (Szczesny *et al*., 2010). *Xe* encodes two T2S systems (designated Xcs- and Xps-T2S), of which only the Xps system contributes to virulence, while the function of the Xcs system remains unclear (Büttner & Bonas, 2010; Szczesny *et al*., 2010; Timilsina *et al*., 2020). The Xps-T2S system is encoded by an eleven-gene chromosomal cluster (*xpsE–M*, *C*, *D*) that is co-expressed with T3S genes, suggesting coordination between these virulence systems (Szczesny *et al*., 2010; Goll *et al*., 2025). T2S-mediated cell wall degradation has been proposed to facilitate assembly of the T3S pilus at the bacterium–plant interface, thereby enhancing T3E delivery into host cells (Szczesny *et al*., 2010).

Previous studies identified five T2Es in *Xe*, three xylanases (XCV0965, XynB2, XynB3), an esterase homolog (LipA), and a serine protease (XCV3671). These were identified based on sequence homology to known T2Es and the presence of Sec or Tat signal peptides (Szczesny *et al*., 2010; Solé *et al*., 2015). Single deletions in these genes reduce *in planta* bacterial growth and disease symptom formation (Tamir-Ariel *et al*., 2012; Solé *et al*., 2015), demonstrating that extracellular proteolytic and cell wall-degrading activities contribute to virulence. However, since no conserved signal for T2S-dependent transport across the OM has been identified, T2E repertoires cannot be predicted computationally (Thomassin *et al*., 2017; Korotkov & Sandkvist, 2019), limiting our ability to identify and characterize these proteins.

In this study, we found that the T2S system from *Xe* substantially promotes bacterial proliferation in tomato independently of T3E delivery. T2Es are required for nutrient acquisition as was shown by bacterial growth assays using plant cell wall polysaccharides as carbon and proteins as nitrogen sources. A similar role was observed for T2Es from *Xanthomonas axonopodis* pv. *glycines* (*Xag*) and *Xanthomonas campestris* pv. *campestris* (*Xcc*), which infect soybean and *Brassica* species, respectively. Quantitative plant cell wall compositional analysis provided direct evidence for T2E-driven cell wall remodeling during infection. Furthermore, *in planta* proteomics identified an arsenal of 22 T2Es from *Xe*, which target all major cell wall polysaccharides. Among the identified T2Es are putative proteases which belong to four distinct catalytic classes and include a novel fungal-like glutamic protease. While different protease repertoires in *Xe*, *Xcc* and *Xag* hint at strain-specific functions, the conservation of the identified CWDEs in all three strains support their predicted core role in cell wall degradation and nutrient acquisition during infection. Together, our findings identify the T2S system as a key nutritional system for bacterial infections of plants.

## Materials and Methods

### Bacterial strains, growth conditions and growth curves

Bacterial strains, growth conditions and growth curves are described in the Supplementary Material and Methods.

### Plant material and plant inoculations

Plant material and inoculations are described in the Supplementary Material and Methods.

### Generation of deletion mutants

*Xanthomonas* mutants were generated via homologous recombination-mediated deletion (Huguet *et al*., 1998). To this end, regions flanking the *xps* gene clusters of *Xe*, *Xag* and *Xcc* strains were amplified from bacterial genomic DNA and cloned in the suicide vector pOGG2 (Goll *et al*., 2025). Plasmids generated are listed in Table S1. *Xanthomonas* strains were transformed and colonies with double crossover events resulting in cluster deletions were identified via PCR with primers flanking the *xps* cluster.

### Callose deposition assays

Analysis of callose deposits was performed as described by Mason *et al*. (2020). Six-mm leaf discs were collected and destained in 100% EtOH for 1 day and subjected to 50% EtOH for 30 min and equilibrated in 67 mM K_2_HPO_4_ (pH 12) for 30 min. Staining solution (0.1% aniline blue solubilized in 67 mM K_2_HPO_4,_ pH 12) was added for 1 h before analysis in mounting solution (70% glycerol, 30 % staining solution). Imaging was performed with an Axio Zoom V16 (Zeiss) equipped with a CFP filter. Callose deposits were quantified automatically using a Fiji macro adapted from Zavaliev and Epel (2015) (Script S1).

### Quantitative reverse transcription (RT)-PCR analysis

For quantitative RT-PCR analysis, Four VF36 tomato plants were inoculated on individual leaflets of the third and fourth leaves with a needleless syringe. Bacteria were suspended in water at OD_600_ of 0.2 (2 × 10^8^ cfu/mL) and flg22 at a concentration of 100 nM. Total RNA was isolated from 6 leaf discs (size 6 mm) per treatment per time point (NucleoSpin RNA plant kit; Macherey-Nagel). 300 ng of total RNA was used for RT (RevertAid First Strand cDNA Synthesis Kit; Thermo Scientific) and then cDNA was diluted 1:4 to use as input in a qPCR reaction using MESA Blue qPCR Mastermix Plus. 7.5 ng of cDNA was used in an 8 µL reaction (3 technical replicates). Primers were designed by Kim *et al*. (2009). For data analysis, Ct values were normalised to ACTIN, and relative expression values were determined against *Xe* WT using the comparative Ct method (2^-ΔΔCt^) (Livak & Schmittgen, 2001).

### Transmission electron microscopy

Pepper leaf samples were fixed in 3% glutaraldehyde in 0.1 M sodium cacodylate buffer (pH 7.2) for 3.5 h, washed three times (10 min each) in buffer, and post-fixed in 1% osmium tetroxide in 0.1 M sodium cacodylate buffer for 1 h. After triple washing in distilled water (10 min each), samples were dehydrated in a graded ethanol series (10%, 30%, 50%, 70%, 90%, and 100%; 30 min each). En bloc staining was performed with 1% uranyl acetate in 70% ethanol for 1 h. Following two washes in 100% ethanol (30 min each), specimens were infiltrated with increasing concentrations of Spurr resin in ethanol (1:3, 1:1, 3:1, and pure Spurr); several hours to overnight per step, then polymerization in fresh Spurr resin at 70°C for 16 h. Ultrathin sections (∼70 nm) were cut with an ultramicrotome (Ultracut S, Leica, Wetzlar, Germany), collected on copper grids and contrasted with uranyl acetate and lead citrate (AC20, Leica). Images were captured using a Zeiss EM900 at 80kV, equipped with a slow scan 1k CCD camera (Tröndle Restlichtverstärkersysteme, Moorenweis, Germany).

### Analysis of cell wall composition and thickness

The analysis of cell wall composition and thickness was performed as is described in the Supplementary Material and Methods.

### Cloning of T2Es

To generate constructs for secretion assays, native promoters (between 200 and 500 bp upstream of the start codon) and coding sequences lacking the stop codon were amplified by PCR from *X. euvesicatoria* (Table S1). Internal *Bsa*I recognition sites were removed using PCR-based mutagenesis. PCR amplicons were sub-cloned into pAGM9121 using *Bpi*I and T4 DNA ligase (Goll *et al*., 2025). Promoters and genes were then assembled into the expression vector pBRM-P using *Bsa*I and T4 DNA ligase (Szczesny *et al*., 2010). For overexpression of enzymes in the BL21 *E.coli* Star strain, coding sequences were ligated into the pBRM expression vector under the control of the constitutive *lac* promoter (Szczesny *et al*., 2010). All coding sequences were cloned in frame with a 3x c-Myc epitope-coding sequence followed by a stop codon to allow detection via western blot.

### Enzyme activity assays

Bacteria were cultured overnight in NYG media (for *Xanthomonas*) or LB media (for *E. coli*) pelleted (8000 ×g) and resuspended to OD_600_ of 1.0 (1 × 10⁹ cfu/mL). Suspensions were pipetted into holes punched in 1% agar plates, with 2% milk, 0.1% remazol brilliant blue (RBB) xylan or 1% CMC to assay for protease, xylanase or cellulase activity, respectively, and incubated at 28°C for 2 days. Bacteria were removed prior to documentation. CMC plates were stained with 0.2% Congo red (Sigma-Aldrich) solution and destained with 0.5 M NaCl to visualize cellulase activity (Gough, 1988).

### Collection of apoplast wash fluid

Four leaflets were collected from third and fourth leaves of three dip-infected Moneymaker tomato plants (4–6 weeks). Midveins were removed and cut edges were rinsed with distilled water. Leaflets were submerged in cold distilled water in a vacuum flask, and infiltrated (O’Leary *et al*., 2014). Leaflet halves were rolled in parafilm and inserted into a 15 mL Eppendorf tube (cut side up) and the parafilm was folded over the tube to hold the leaves in place. Samples were centrifuged at 250×g for 15 min at 4°C and apoplast wash fluid (APW) was collected from the bottom of tube. Fluid collected from leaflets was pooled by plant. Samples were centrifuged at 15,000×g for 5 min at 4°C to remove cell debris. Apoplast wash fluid samples were prepared as described in Supplementary Material and Methods.

### Secretion assays

For *in vitro* secretion assays, 15 mL of XVM2 medium were inoculated from an NYG overnight liquid culture with a bacterial density of 2×10^7^ cfu/mL and incubated at 30°C for 3-4 h shaking at 200 rpm. Total extracts and supernatants were collected as described previously (Rossier et al., 1999), and equal amounts adjusted according to the bacterial density were analysed by SDS-PAGE and immunoblotting, using a c-Myc epitope-specific antibody (polyclonal antibody from rabbit, Sigma-Aldrich).

## Results

### The T2S system promotes growth of *Xe* in tomato independently of the T3S system

Our previous studies suggested a contribution of T2Es in the assembly of the T3S pilus across the plant cell wall (Szczesny *et al*., 2010). To test whether T2S system-mediated virulence is solely the result of reduced T3S system functionality during infections, we analysed the combined and individual contributions of the T2S and T3S systems to the interaction in the *Xe* (strain 85-10)*-*tomato pathosystem. For this, the *in planta* growth of bacterial mutants with deletions of the T3S system ATPase HrcN (Δ*hrcN*), the entire *xps* gene cluster (Δ*xps*) and both systems (*ΔxpsΔhrcN*) were analysed in tomato. When infiltrated into leaves of susceptible tomato plants, the Δ*xps* mutant strain grew significantly less than the wild-type strain starting at 5 days post-infection (dpi), a trend that persisted through 16 dpi. This demonstrated the importance of the T2S system to *Xe* proliferation in tomato (Fig. 1a). The Δ*hrcN* mutant exhibited a comparable replication deficit (Fig. 1a). Consistent with reduced bacterial proliferation, bacterial spots were not visible following *Δxps* mutant or *ΔhrcN* mutant inoculations of tomato leaves (Fig. 1b). The pronounced reduction in bacterial growth of *Δxps* mutant strain, on parallel with that of the Δ*hrcN* mutant, was unexpected given that the T3S system is considered the primary virulence determinant of *Xe* (Büttner & Bonas, 2010). The double mutant strain displayed an even greater reduction in bacterial titers, indicating that both secretion systems contribute additively to full virulence. This suggests that beyond any supportive role the T2S system may play in T3S system function, it also fulfills distinct, independent functions essential for tomato colonization.

**Fig. 1.**
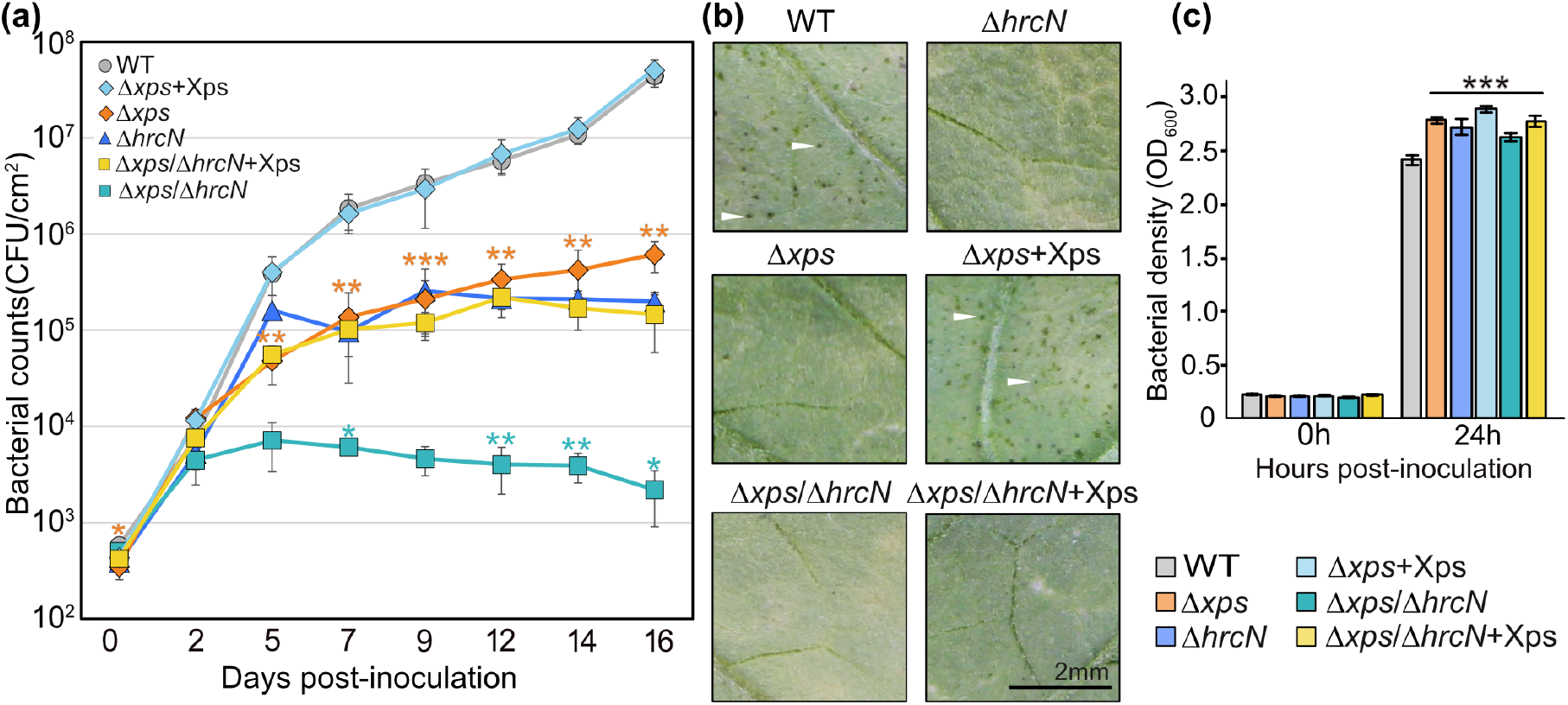
The T2S system functions independently of the T3S system. Money Maker tomato leaves were syringe-inoculated with 5×10⁴ cfu/mL of *Xe* wild type strain 85-10 (WT, gray), the T2S system mutant (Δ*xps*, orange), T3S system mutant (Δ*hrcN*, in dark blue), the double mutant (Δ*xps*/Δ*hrcN*, turquoise) and the complemented T2S system mutant and double mutant expressing the modular *xps*-T2S system on a plasmid (Δxps+Xps in light blue and *Δxps/ΔhrcN*+Xps in yellow, respectively). **(a)** *In planta* bacterial multiplication over 16 days expressed as mean cfu/cm^2^ (n = 4 plants). Experiments were repeated 2 times. Error bars are standard deviation. Significant Welch’s t-test results between Δ*xps* and WT or Δ*hrcN* and the double mutant are indicated with asterisks: *p < 0.05, **p < 0.01, ***p < 0.001. Δ*xps*, Δ*hrcN* and Δ*xps*/Δ*hrcN*+Xps were not significantly different. **(b)** Plant phenotypes at 22 days post-inoculation. Examples of water-soaked lesions are marked with white arrows. The scale bar applies to all images. **(c)** Bacterial growth in liquid NYG culture medium at 0 and 24 h expressed as mean OD_600_ values (n = 3). Experiments were repeated three times. Error bars are standard deviations. One-way ANOVA followed by Dunnett’s multiple comparison test was performed compared to wild type: ***p < 0.001.

Complementation with the *xps*-T2S system gene cluster (+Xps) on a plasmid restored *in planta* bacterial growth of both Δ*xps* and double mutant (Δ*xps*Δ*hrcN*) strains to wild-type and Δ*hrcN* mutant levels, respectively, confirming that the growth defects were T2S system-specific (Fig. 1a). Mutant growth in culture was comparable across strains, ruling out general multiplication defects (Fig. 1c).

These findings demonstrate that the T2S system is a key determinant of *Xe* virulence in tomato, even when the T3S system is inactive. We therefore investigated whether the T2S system contributes to (1) nutrient acquisition through plant cell wall degradation, and (2) PTI suppression.

### *Xe* strains with a functional T2S system elicit PTI responses in tomato

To examine whether the virulence function of the T2S system in tomato is linked to the suppression of plant defense responses, we analysed callose deposition in infected VF36 tomato leaves. VF36 plants were easier to inoculate than MoneyMaker, while disease symptoms following infection with *Xe* were comparable in both cultivars (Fig. S1). Callose accumulation in leaves is widely used as a readout for PTI responses and can be triggered by recognition of T2Es or by DAMPs released during T2E-mediated cell wall degradation (Hou *et al*., 2019; Lorrai & Ferrari, 2021; Wang *et al*., 2021; Saberi Riseh *et al*., 2024). Tomato leaves were inoculated with the *Xe* wildtype, the Δ*xps*, Δ*hrcN* and double mutant strains and a water control, and callose deposition was measured 24 h post-inoculation (hpi) (Fig. 2). At this time point, bacterial population density *in planta* was comparable across strains (Fig. 2b). Infection with the wild-type strain, the Δ*xps* and the Δ*xps*Δ*hrcN* double mutant resulted in minimal callose deposition at 24 hpi, with levels comparable to those observed in water-treated plants. In contrast to the Δ*xps*Δ*hrcN* double mutant strain, the Δ*hrcN* deletion mutant induced a significant increase in callose deposition, suggesting that collectively T2Es or the T2S system elicit PTI, rather than suppressing it, when the T3S system is inactive (Fig. 2a and 2c). We do not exclude the possibility that individual T2Es may have roles in immune suppression. These findings also suggest that T3Es are critical for suppressing T2S-induced plant immune responses.

**Fig. 2.**
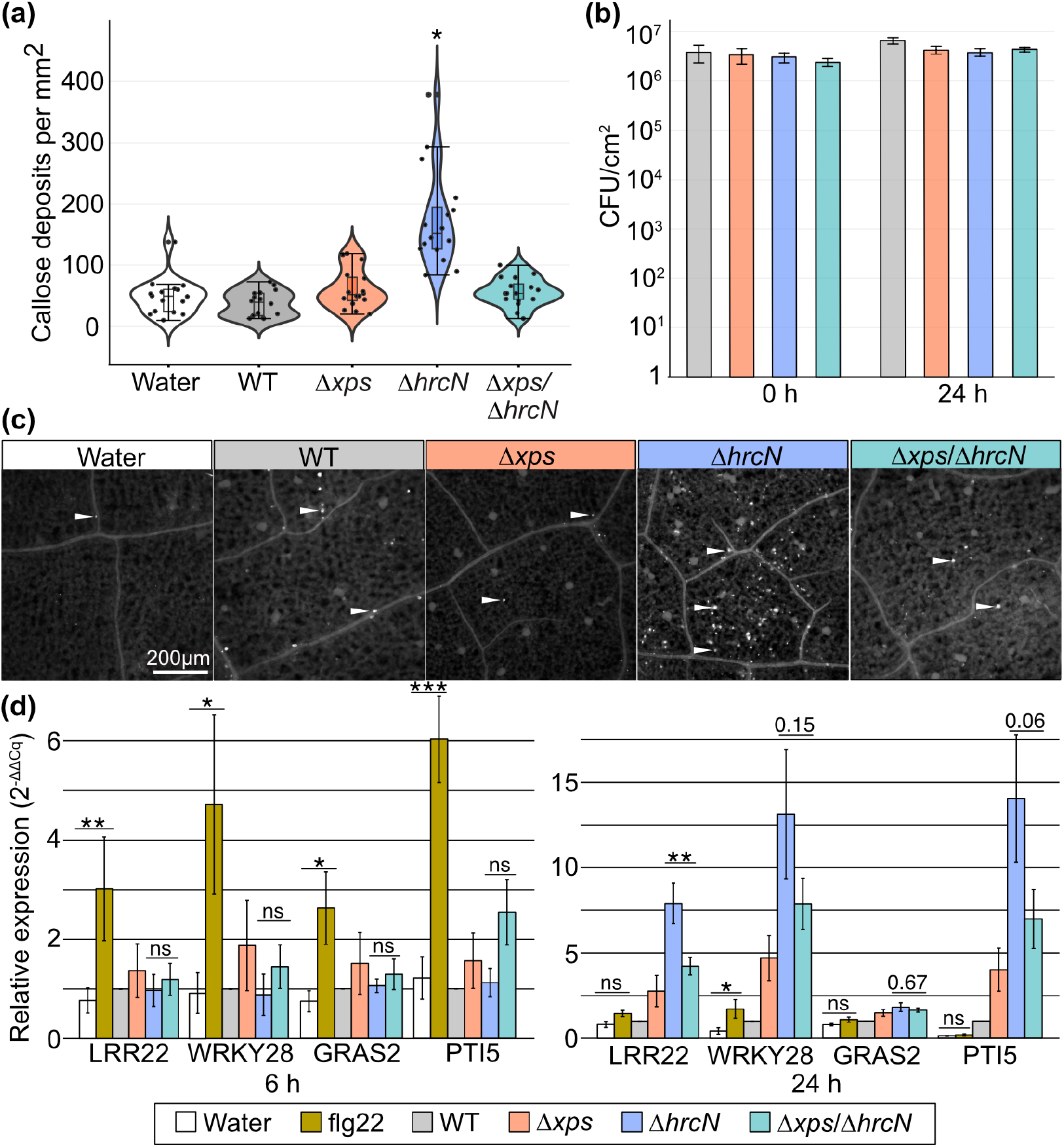
*Xe* T3Es suppress callose deposition triggered by T2E secretion. VF36 tomatoes leaves were infiltrated with water or *Xe* wild-type (WT), T2S system mutant (Δxps, orange), T3S system mutant (Δ*hrcN*, blue) or the double mutant (Δ*xps*/Δ*hrcN*, turquoise) at an OD_600_ of 0.2 (2 × 10^8^ cfu/mL). Callose deposits were quantified 24 hours post infection. **(a)** Quantification of callose deposits per mm^2^. Results are presented as violin plots displaying the median, interquartile range, and data distribution. Due to non-normal distribution (Shapiro-Wilk test), data were transformed using a square root transformation prior to analysis. Differences between treatments were assessed using a one-way ANOVA and Tukey’s post-hoc test. An asterisk denotes a significant difference between treatments. **(b)** Bacterial growth in infected tissue showed no significant differences between strains (same color coding as in a). Error bars are standard deviations. **(c)** Fluorescence microscopy images (CFP filter) of callose deposits, visible as white spots (examples marked with white arrows). The scale bar applies to all images. **(d)** Quantitative RT-PCR of PTI-responsive genes at 6 and 24 h post-infiltration, using the same strains and experimental setup as in callose deposition assays. flg22 peptide suspended in water (100 nM) was added as a positive control for PTI marker gene induction. ACTIN expression was used for normalisation and relative expression was calculated against *Xe* WT using the comparative Ct method (2^-ΔΔCt^). Error bars show the standard error of mean relative expression values (n = 4 plants). Paired t-tests are shown for the flg22 vs water control and Δ*hrcN* vs Δ*xps*/Δ*hrcN* samples, with p-values adjusted using the Holm method. The experiment was repeated twice with similar results.

To examine the dynamics of immune induction by T2Es, we also checked the expression of early PTI marker genes at 6 h and 24 h. We chose four genes previously shown to be responsive during infection with the *Xe* T3S mutant (*ΔhrpF*): *PTI5* (ethylene-responsive transcription factor), *GRAS2* (biotic stress-responsive transcription factor), *WRKY28* (biotic stress-induced transcription factor) and *LRR22* (receptor-like kinase) (Navarro *et al*., 2004; Kim *et al*., 2009). Purified flg22 peptide, a derivative of bacterial flagellin which elicits PTI, was used as a positive control for early immune induction (Zipfel et al., 2004). At 6 h, treatment with flg22 clearly and significantly induced the expression of all four PTI marker genes, a response that diminished after 24 h (Fig. 2d). By comparison, T2E-dependent PTI marker gene induction occurred at a later time point. *LRR22*, *WRKY28* and *PTI5* showed elevated expression levels during infection with the Δ*hrcN* mutant than with the Δ*xps*Δ*hrcN* double mutant, although the difference was statistically significant only for *LRR22*. These results are consistent with the callose deposition data, suggesting that the Δ*hrcN* mutant triggers plant immune responses more effectively than the double mutant. Also consistent with callose deposition, inoculation with wild-type *Xe* consistently resulted in low marker gene expression (p>0.05) at 24 hours compared to Δ*hrcN* inoculations, suggesting that PTI marker gene expression is effectively suppressed by the T3S system. The exception was GRAS, which showed no significant differences between treatments.

### The T2S system is an important virulence factor of xanthomonads

Given the universal presence of the T2S system across all sequenced *Xanthomonas* strains (Alvarez-Martinez *et al*., 2021), we hypothesized that T2Es may serve a fundamental role in nutrient acquisition. To test this, we examined its importance across evolutionarily distant pathosystems. We first tested the mesophyllic *Xag* 8ra, which is closely related to *Xe* and causes bacterial pustule disease on soybean (*Glycine max*). Since *Rosidae* (soybean) and *Asteridae* (tomato and pepper) diverged over 120 million years ago (Magallon *et al*., 2015), they allowed us to test whether the T2S system contributes to bacterial virulence in two distinct dicotyledonous plant lineages. T2S system mutants were generated in *Xag* via deletion of the entire *xps-*T2S gene cluster. For infection studies, soybean leaves were infiltrated with the wild-type *Xag* strain, which led to the expected chlorotic and necrotic disease symptoms (Kim *et al*., 2003). In contrast, the Δ*xps* mutant showed reduced symptoms, suggesting that the T2S system contributes to virulence of *Xag* in soybean (Fig. 3a). Interestingly, introduction of the *xps-*T2S gene cluster from *Xe* into the *Xag* Δ*xps* mutant restored the wild-type phenotype (Fig. 3a).

**Fig. 3.**
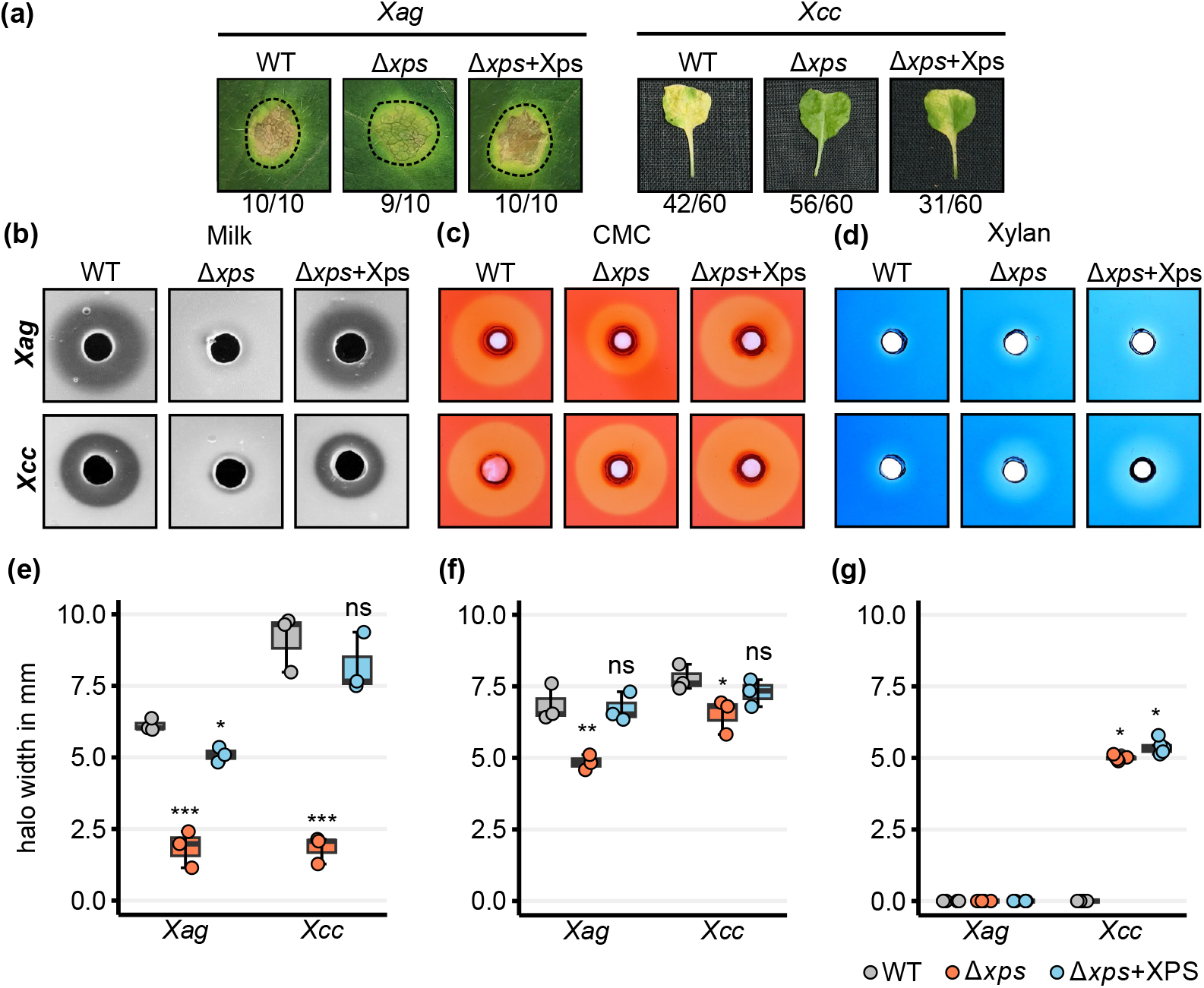
The T2S system from *Xe* complements *Xag* and *Xcc* mutant phenotypes during infections and in plate assays. **(a)** Symptoms following syringe inoculations of soybean (*Glycine max*) with *Xag* (OD_600_ of 0.02; 2×10^7^ cfu/mL) and following clip inoculation of *A. thaliana* (Oy-0) with *Xcc* (OD_600_ of 0.1). Plant were inoculated with wild-type (WT) (Barger *et al*.), Δ*xps* mutant and complementation strains carrying the *xps*-T2S gene cluster from *Xe* on a plasmid (Δ*xps*+Xps). Dashed lines denote inoculated areas on soybean. The number of times phenotypes were observed out of the total inoculations is noted below each image. **(b)** Extracellular protease activity was assayed on 2% milk. **(c)** Extracellular cellulase activity was documented on CMC agar plates where halos were visualized via CMC staining with Congo red. **(d)** Xylanase activity was monitored on RBB-xylan agar plates. All plate assays were incubated for 2 days at 28°C. (**e-g)** Halo size quantification. Box plots display the median (center line), interquartile range (box) and 1.5× interquartile range (whiskers). All data points for a single experiment are shown. One-way ANOVA followed by Dunnett’s multiple comparison test was performed to compare mutants and complementation strains to WT (*p<0.05, **p < 0.01, ***p < 0.001, ns = not significant). Error bars are standard deviations. Experiments were repeated 3 times with similar results.

The T2S system was also described as virulence determinant of *Xcc* (Qian *et al*., 2005; Paauw *et al*., 2024), which is distantly related to *Xe* and *Xag.* While *Xe* and *Xag* are both mesophyll dwelling pathogens, *Xcc* has a vascular and systemic lifestyle (Büttner & Bonas, 2010; Ryan *et al*., 2011). To test the importance of the T2S system in *Xcc*, we generated a deletion mutant lacking the *xps* gene cluster. This was done in an Δ*avrAC* mutant background, as this T3E restricts host range by inducing ETI in some Arabidopsis accessions (Xu *et al*., 2008). Clip infections of Arabidopsis (Oy-0) with the Δ*avrAC* mutant led to visible yellowing of whole leaves indicative of systemic spread throughout the leaf as was described by Paauw *et al*. (2024). The Δ*avrAC*Δ*xps* mutant showed reduced to almost no yellowing (Fig. 3a). Notably, the wild-type phenotype was restored in the *Xcc* Δ*xps* mutant via introduction of the *Xe xps* gene cluster. The conservation of *xps* cluster proteins across strains (Table S2) is consistent with the functional interchangeability demonstrated by heterologous complementation.

### The Xps-T2S system contributes to extracellular protease and cellulase activity of *Xag* and Xcc

Building on our previous characterization of *Xe* T2E activities (Szczesny *et al*., 2010), we tested extracellular enzyme activities of *Xcc* and *Xag* Δ*xps* mutants on milk-, carboxymethylcellulose (CMC)- and xylan-containing agar plates. Here, halo formation indicates substrate degradation by secreted enzymes. *Xag* and *Xcc* Δ*xps* strains showed reduced milk protein degradation compared to the respective wild-type strains, suggesting that the secretion of proteases is dependent on the T2SS (Fig. 3b). Deletion of the *xps-*T2S gene cluster reduced but did not abolish extracellular cellulase activity, revealing that some cellulases are secreted via the T2S system (Fig. 3c). The biggest differences were observed in xylan utilization. *Xag* and *Xcc* activity on RBB-xylan was very minimal. Halos produced by *Xcc* Δ*xps* actually increased in size compared to the wild-type (Fig. 3d). Protease and cellulase activities were restored by complementation with the *Xe xps* gene cluster (Fig. 3d-g).

### The T2S system promotes bacterial growth on plant cell wall extracts

To investigate a role of the T2S system from *Xe, Xag* and *Xcc* in nutrient acquisition through cell wall degradation, strains were cultivated in modified minimal medium (XVM2), in which sucrose and fructose were replaced by isolated leaf cell wall extracts (alcohol-insoluble residue, AIR) from tomato, pepper, soybean or Arabidopsis as sole carbohydrate source. While the monosaccharide composition of all four species was dominated by galacturonic acid and cellulose, each species displayed a distinct compositional profile (Table S3). Wild-type *Xanthomonas* strains grew on all four plant cell wall extracts, suggesting their capacity to utilize cell wall components from a range of dicot hosts (Fig. 4). In all cases, *xps* mutants showed reduced growth, highlighting the importance of the T2S system for metabolizing plant cell wall components in nutrient-limiting conditions. Ectopic expression of the *Xe xps* T2S gene cluster restored growth of all *xps* mutants to near wild-type levels (Fig. 4). It should be noted that AIR preparations likely retain cell wall-associated proteins in addition to structural polysaccharides. Therefore, the observed differences between strains likely reflect differences in their ability to utilize plant cell wall material as a whole, including its associated components.

**Fig. 4.**
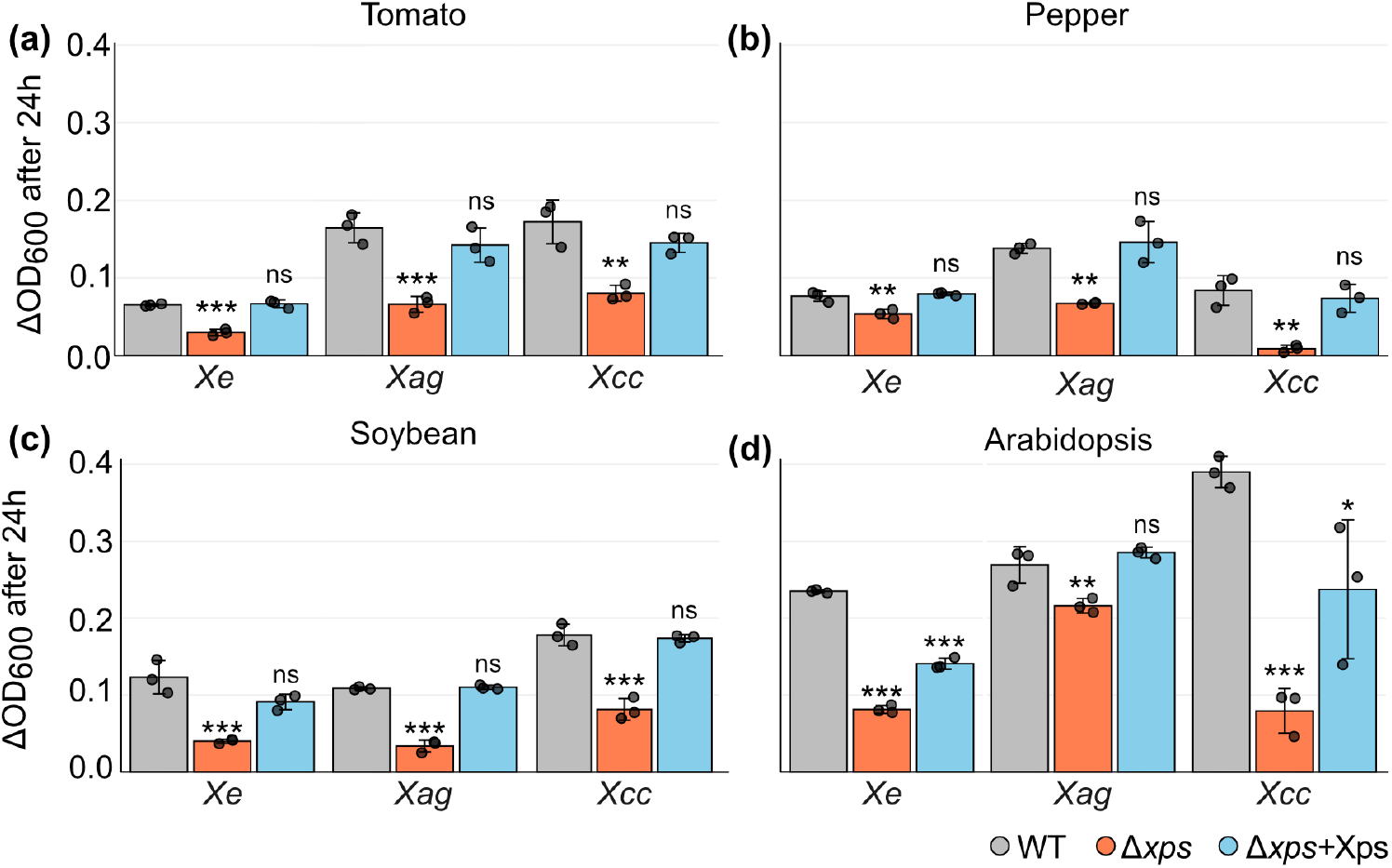
T2S system-dependent growth of *Xanthomonas* on cell wall extracts of diverse dicot species. Wild-type (WT, gray) *Xe*, *Xag* and *Xcc* strains, corresponding T2S mutants (Δ*xps*, orange) and strains complemented with the modular *xps* T2S gene cluster (Δ*xps*+Xps, blue) were grown in XVM2 medium (without fructose or sucrose), which was supplemented with 0.5 mg/mL cell wall extracts from **(a)** tomato, **(b)** pepper, **(c)** soybean or **(d)** Arabidopsis leaves. Growth is expressed as mean change in culture optical density (ΔOD_600_) after 24 h (n=3). All cultures we started with an OD_600_ of 0.02 (2×10^7^ cfu/mL). Dots represent individual data points. Experiments were repeated 3 times with similar results. Data normality was assessed using Shapiro-Wilk tests and variance homogeneity via Levene’s test. One-way ANOVA followed by Dunnett’s multiple comparison test was performed. **p < 0.01, ***p < 0.001, ns = not significant. Error bars are standard deviations.

### T2Es contribute to the metabolization of plant cell wall components

To specifically investigate T2E contributions to the metabolism of cell wall polysaccharides we made use of commercially available CMC (modified cellulose), xylan (from corn cob) and pectin (from citrus peel). We performed growth assays in XVM2 medium supplemented with glucose (positive control), CMC, pectin or xylan as sole carbohydrate sources. All strains grew on all substrates (Fig. 5), though growth of *Xe* was minimal on CMC compared to *Xag* and *Xcc*. All strains multiplied poorly on pectin. This was unexpected, given that xanthomonads encode predicted pectin-modifying enzymes (Kaewnum *et al*., 2006; Wang *et al*., 2008; Tayi *et al*., 2016). Bacteria grew well on xylan (Fig. 5d), fitting with the identification of xylan metabolism genes present in xanthomonads (Dejean *et al*., 2013; Vieira *et al*., 2021).

**Fig. 5.**
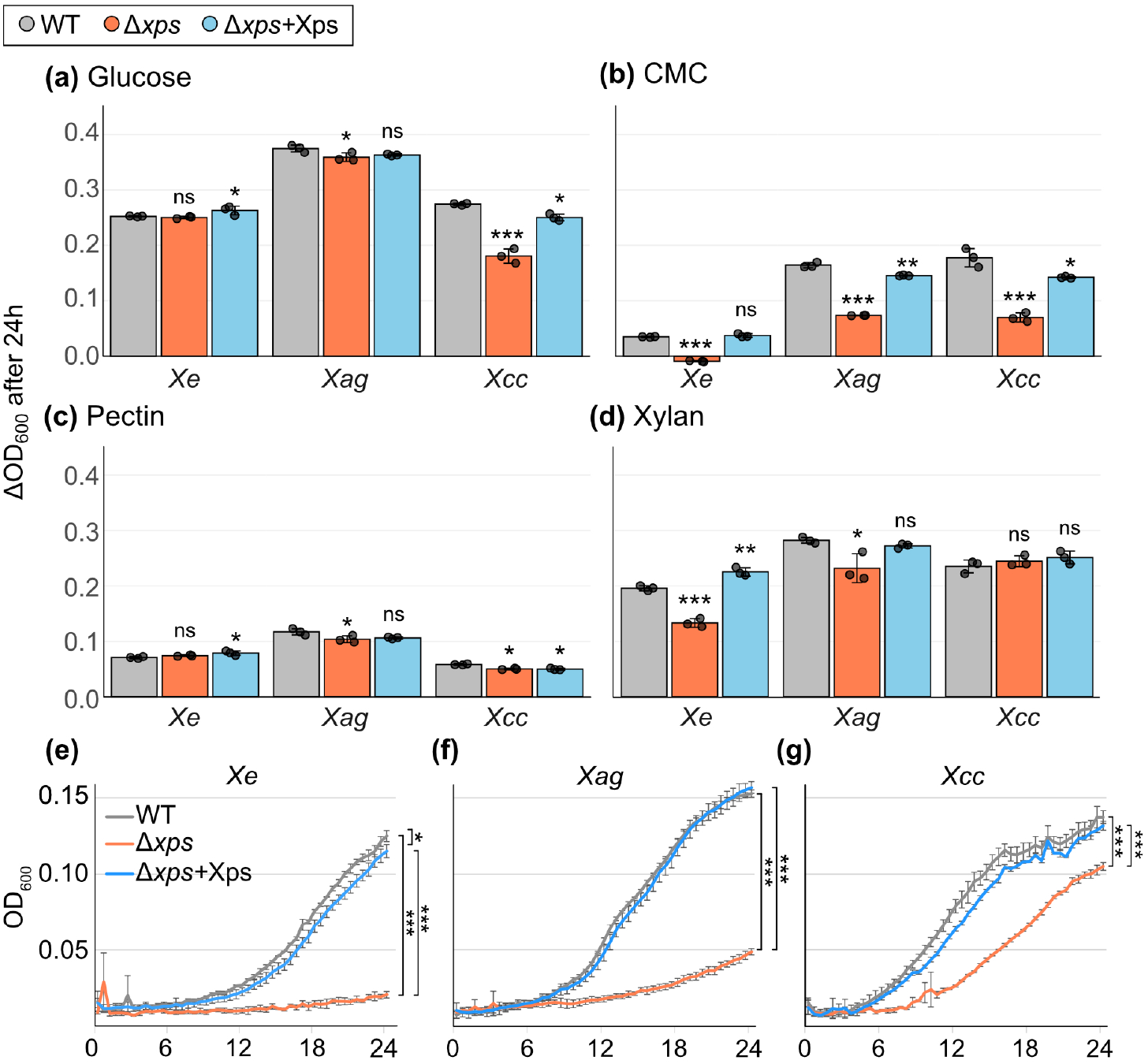
*Xanthomonas* metabolizes commercially purified cell wall components and milk proteins. Wild-type (WT, gray) *Xe*, *Xag* and *Xcc*, T2S system mutants (Δ*xps*, orange) and strains complemented with the modular *Xe xps* T2S gene cluster (Δ*xps*+Xps, blue) grown in XVM2 medium (without fructose and sucrose) supplemented with commercial cell wall components as a carbon source: **(a)** 0.4 mg/mL glucose **(b)** 4 mg/mL carboxymethylcellulose (CMC), **(c)** 0.4 mg/mL pectin (from citrus peel), or **(d)** 0.4 mg/mL xylan (from corn cob). Growth is expressed as mean change in culture optical density (ΔOD_600_) after 24 h (n = 3). All cultures we started with an OD_600_ of 0.02 (2×10^7^ cfu/mL). Dots represent individual data points Experiments were repeated 3 times with similar results. Error bars represent standard deviation. Data normality and variance homogeneity were assessed using Shapiro-Wilk tests and Levene’s test, respectively. For normally distributed data, one-way ANOVA followed by Dunnett’s multiple comparison test was performed. For non-normally distributed data, Kruskal-Wallis test followed by Conover post-hoc test with Holm-Šídák correction was used. Datasets for *Xag* in CMC-, *Xcc* in pectin- and *Xe* in glucose-containing medium were analysed using Kruskal-Wallis tests; for all other experiments ANOVA was used (*p < 0.05, **p < 0.01, ***p < 0.001, ns = not significant). Growth of **(e)** *Xe*, **(f)** *Xag* and **(g)** *Xcc* WT, Δ*xps* and strains complemented with the *Xe xps* gene cluster (+Xps) was analysed in liquid XVM2 medium containing 0.03% skim milk powder as the sole amino acid source. Cultures were inoculated with an OD_600_ of 0.02 and monitored for 24 h. Statistically significant differences were determined at the 24 h timepoint using one-way ANOVAs with post-hoc Tukey-HSD (*p<0.05, ***p<0.001). Experiments were performed at least three times; one representative dataset is shown.

The role of the T2S system varied across strains. In *Xag*, the T2S system contributed to growth on all substrates including glucose. The *Xcc* Δ*xps* mutant multiplied less than the wild-type strain on cellulose and glucose but not on pectin or xylan. In *Xe*, the T2S system was required for growth on cellulose and xylan but not on glucose or pectin. These differences suggests that *Xe*, *Xcc* and *Xag* vary in their T2E repertoires. In contrast, components of the T2S machinery likely perform conserved functions across these strains, as demonstrated by the results of our complementation studies with the *Xe xps* gene cluster.

Given that proteins are also an essential part of plant cell walls (Tan *et al*., 2013), and T2S system-dependent protease secretion is reliably observed (Fig. 3), we also evaluated a possible role of proteolysis in nutrient acquisition. Therefore, we cultivated wild-type strains and *xps* mutants of *Xe*, *Xag*, and *Xcc* in modified XVM2 medium in which free amino acids were substituted with skim milk powder. All three wild-type strains multiplied under these conditions (Fig. 5e-g), whereas growth of *Xe* and *Xag xps* mutants was severely impaired, indicating the critical requirement for T2S system-mediated efficient protein utilization (Fig. 5e and 5f). In contrast, the *Xcc* Δ*xps* mutant grew substantially better, suggesting reduced reliance on the T2S system (Fig. 5g). Growth was restored by the *Xe xps* gene cluster in all strains. Together this indicates that protein breakdown contributes to bacterial nutrition alongside carbohydrate degradation.

### *Xe* infection alters the cell wall composition in infected pepper leaves

To assess T2E-driven cell wall remodeling *in planta*, we determined the cell wall monosaccharide composition in pepper leaves which were syringe-inoculated with *Xe* wild-type, Δ*xps* mutant and the complementation strain or MgCl_2_ as a control. Samples were taken 5 dpi, when differences between bacterial growth are minimal (Fig. 6a). Importantly, AIR samples from *Xe*-infected leaves contained monosaccharides of bacterial and plant origin. Bacterial cell walls are composed of peptidoglycan, which consists of N-acetylglucosamine, N-acetylmuramic acid and lipopolysaccharides (Volk, 1966; Boulanger *et al*., 2014; Ogasawara & Dairi, 2021). Accordingly, analysis of cultured *Xe* detected glucosamine, the hydrolysis product of N-acetylglucosamine, mannose, rhamnose and high amounts of glucose (Table S4). In addition, bacteria also secrete xanthan, which consists of glucose, mannose and glucuronic acid (Bhat *et al*., 2022). Mannose and glucuronic acid, which are relatively rare in plant cell walls (Delmer *et al*., 2024; Table S3), were strongly abundant in AIR samples from infected leaves due to the presence of bacteria (Fig. 6b).

**Fig. 6.**
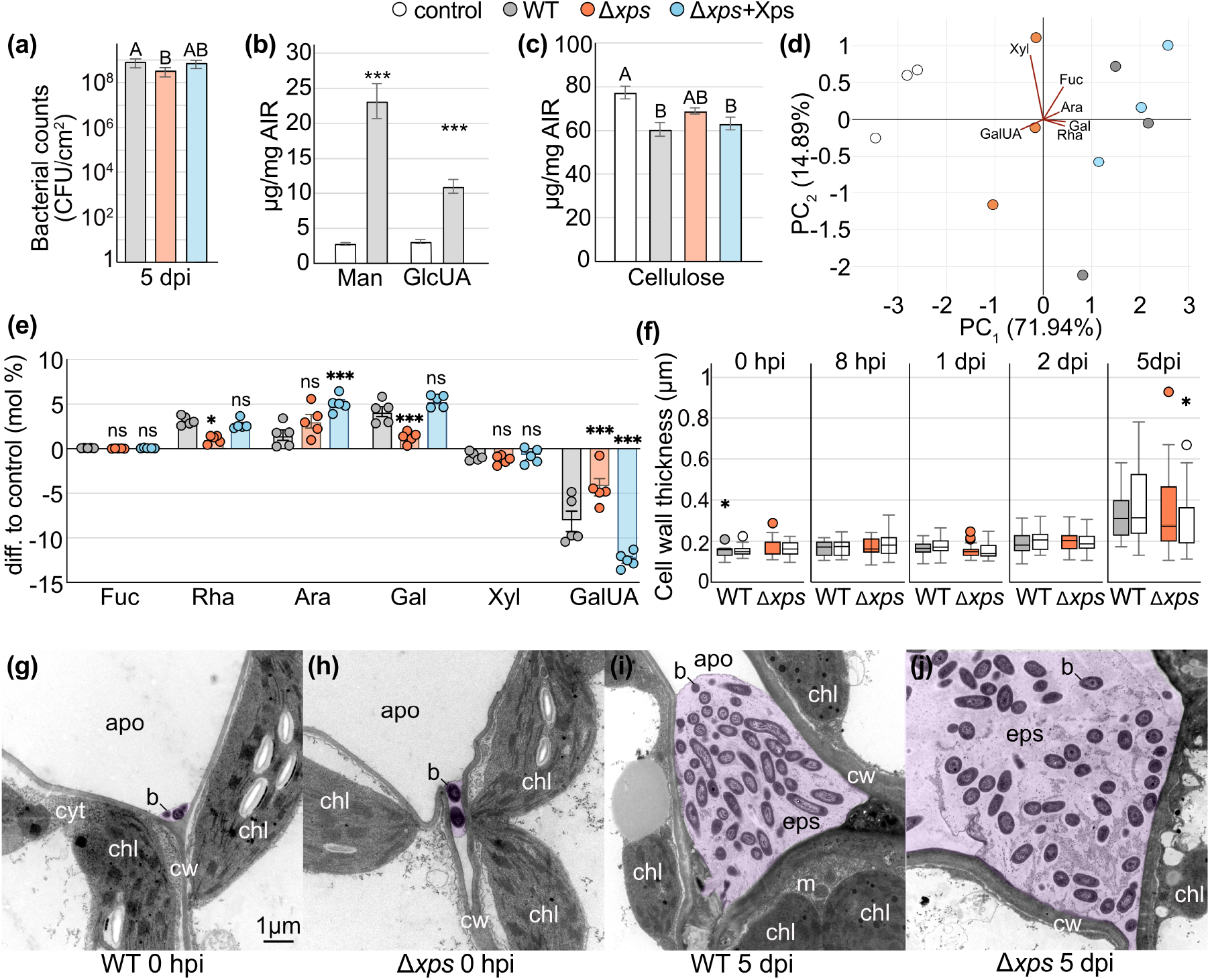
T2S-dependent changes in pepper cell wall composition during infection. Pepper plants were inoculated with wild-type Xe (WT, gray), T2S system mutants (Δ*xps*, orange), or strains complemented with the modular *Xe xps* T2S gene cluster (Δ*xps*+Xps, blue) via needless syringe at a bacterial density of 2 × 10^7^ cfu/mL. MgCl_2_ was infiltrated as a control (white). **(a)** Bacterial counts at 5 dpi from tissues used for the cell wall analysis. Treatments that were significantly different according to a one-way ANOVA and Tukey’s post-hoc were marked with different letters. **(b)** Mannose (Man) and glucuronic acid (GlcUA) amounts in alcohol-insoluble residue (AIR) of pepper leaves were quantified at 5 dpi with *Xanthomonas euvesicatoria*. Values are means of 5 biological replicates and error bars represent the standard error of the mean. Asterisks indicate statistically significant differences to control treatments according to two-way ANOVA and Holm-Šídák’s test (***p < 0.001). **(c)** Cellulose abundance in AIR of pepper leaves at 5 dpi after infection with *Xe* WT, Δ*xps* and Δ*xps*+Xps compared to MgCl_2_-inoculated control (N=5, ± standard error of the mean). Different letters indicate statistically significant differences to control treatment according to one-way ANOVA and Holm-Šídák’s test (p < 0.05). **(d)** Principal component analysis of plant-specific monosaccharides at 5 dpi after infection with *Xe* WT, Δ*xps* and Δ*xps*+Xps compared to the mock control. Data represent mean values from three independent experiments. Xyl, xylose; Fuc, fucose; Ara, arabinose; Gal, galactose; Rha, rhamnose; GalUA, galacturonic acid. **(e)** Relative changes in Fuc, Rha, Ara, Gal, Xyl, and GalUA plant cell wall monosaccharides of infected tissues expressed as mol% difference to MgCl_2_ control (N = 5 ± standard error of the mean). Asterisks indicate statistically significant differences to Xe WT according to two-way ANOVA and Dunnett’s test (*p < 0.05, **p < 0.01, ***p < 0.001, ns = not significant). **(f)** Measurements of cell wall thickness taken at different time points after infection with *Xe* WT and the Δ*xps* mutant taken from areas where bacteria were in contact with cell walls (colored boxes) and bacteria-free areas (white boxes). Box plots show the median (center line), interquartile range (IQR; box) and whiskers extend to 1.5x IQR. Outliers are plotted as individual points. Statistically significant differences between control regions and regions in contact with bacteria according to Mann-Whitney U tests are denoted with an asterisk (*p < 0.05). **(g)-(j)** Electron microscopy images of pepper leaves infected with *Xe* WT or the Δ*xps* mutant at 0 and 5 dpi. apo = apoplast, b = bacteria, chl = chloroplast, cw = cell wall, eps = extracellular polysaccharides (false-colored purple), m = mitochondria.

Plant cell walls are composed of three major classes of polysaccharides: cellulose, hemicellulose, and pectin, each contributing a distinct set of monosaccharides. As cellulose is more resistant to acid hydrolysis than the other classes, it can be precisely isolated and quantified. Compared with MgCl_2_-infiltrated controls, infection with the *Xe* wild-type strain significantly reduced the cellulose content, an effect that was slightly less pronounced after infection with the Δ*xps* mutant (Fig. 6c). Hemicelluloses and pectin together form an interconnected cell wall matrix, which comprises the monosaccharides glucose, fucose, rhamnose, arabinose, galactose, xylose, mannose and uronic acids (Delmer *et al*., 2024). To investigate if monosaccharide composition of the complex polysaccharides present in hemicellulose and pectin depends on the T2S system, we performed principal component analysis (PCA) of non-cellulosic plant cell wall matrix monosaccharides obtained from three independent experiments. We excluded monosaccharides with high abundance in *Xe* cell walls or xanthan from the analysis (glucose, mannose, and glucuronic acid) and investigated quantitative data of fucose, rhamnose, arabinose, galactose, xylose, and galacturonic acid. PC_1_, explaining 71.94 % of the variation in the dataset, clearly separated control, Δ*xps*- and wild-type-infected samples, and was mainly influenced by galacturonic acid, arabinose, galactose and rhamnose (Fig. 6d). Instead, samples infected with the complemented strain clustered together with wild-type-infected samples demonstrating T2S-dependent compositional changes detectable across whole-leaf tissue (Fig. 6d). To Investigate T2S-dependent shifts in more detail, we calculated the relative changes in the monosaccharide composition in one representative experiment. Compared with control plants, infection with wild-type *Xe* led to mild reduction in the relative level of xylose and strong reduction in the relative level of galacturonic acid, respectively, accompanied by increases in the relative levels of rhamnose, arabinose and galactose (Fig. 6e, Table S5). The changes in rhamnose, galactose and galacturonic acid were significantly attenuated in leaves infected with the Δ*xps* mutant, but not with the complemented line, highlighting specific T2S-dependent changes *in planta* (Fig. 6e, Table S5). The observed depletion of galacturonic acid from cell walls suggests that pectinases are active *in planta* despite the apparent inability of the bacteria to metabolize commercial pectin *in vitro* and may therefore function as virulence factors independently of bacterial nutrition.

When analysed by electron microscopy, infected leaf tissue did not show T2S-dependent modifications of the plant cell wall ultrastructure (Fig. 6f-j). Measurements of cell wall thickness revealed no significant differences at any time point tested (Fig. 6f, Table S6). Although post-infection thickening was observed at 5 dpi, it occurred in all treatments. Imaging showed rapid colonization of intercellular spaces, with bacteria embedded in an EPS matrix by 5 dpi and few cells were in direct contact with plant cell walls (Fig. 6g-j).

### Identification of T2Es from *Xe* in the apoplastic fluid of infected tomato plants

The identity of T2Es from *Xe,* which contribute to cell wall degradation and bacterial nutrition, remained to be investigated. Previously, the predicted xylanases XCV0965, XCV4358 and XCV4360, the putative serine protease XCV3671 and the lipase LipA (XCV0536) were identified as T2Es through direct testing of candidate proteins in *in vitro* secretion assays (Szczesny *et al*., 2010; Tamir-Ariel *et al*., 2012; Solé *et al*., 2015). However, since no conserved signal for T2S system OM transport has been identified, computation prediction of T2E repertoires is not possible (Thomassin *et al*., 2017; Korotkov & Sandkvist, 2019). The abundance of predicted carbohydrate-active enzymes in the genome of *Xe* (Table S8) emphasizes the importance of an unbiased approach to identify secreted T2Es from the plant apoplast during infection.

Tomato plants were dip-infected with *Xe* wild-type or Δ*xps* mutant strain and apoplast wash fluid (APW) was collected at 4 dpi and analysed via label-free proteomics. Mass spectrometry identified 87 Xe proteins (Table S7), of which 24 were consistently less abundant in APW from Δ*xps*-infected plants, suggesting T2S-dependent secretion (Fig. 7a). Two were excluded due to ambiguous peptide assignment. Multiple unique peptides were detected per protein (5–23 peptides; 13.7–74.5% sequence coverage), and all except the adhesin XadA2 contained predicted signal peptides, which mediate transport into the periplasm (Signal P 6.0 prediction, Teufel *et al*. (2022)). Recovery of known T2Es XCV4358 and XCV3671 validated the approach. OM proteins (TonB receptors, OmpA, OmpW3 and HrpF), which are typical of OM vesicles (Sidhu *et al*., 2008), were enriched in samples infected with the Δ*xps* mutant, suggesting a possible increase in the absence of T2S.

**Fig. 7.**
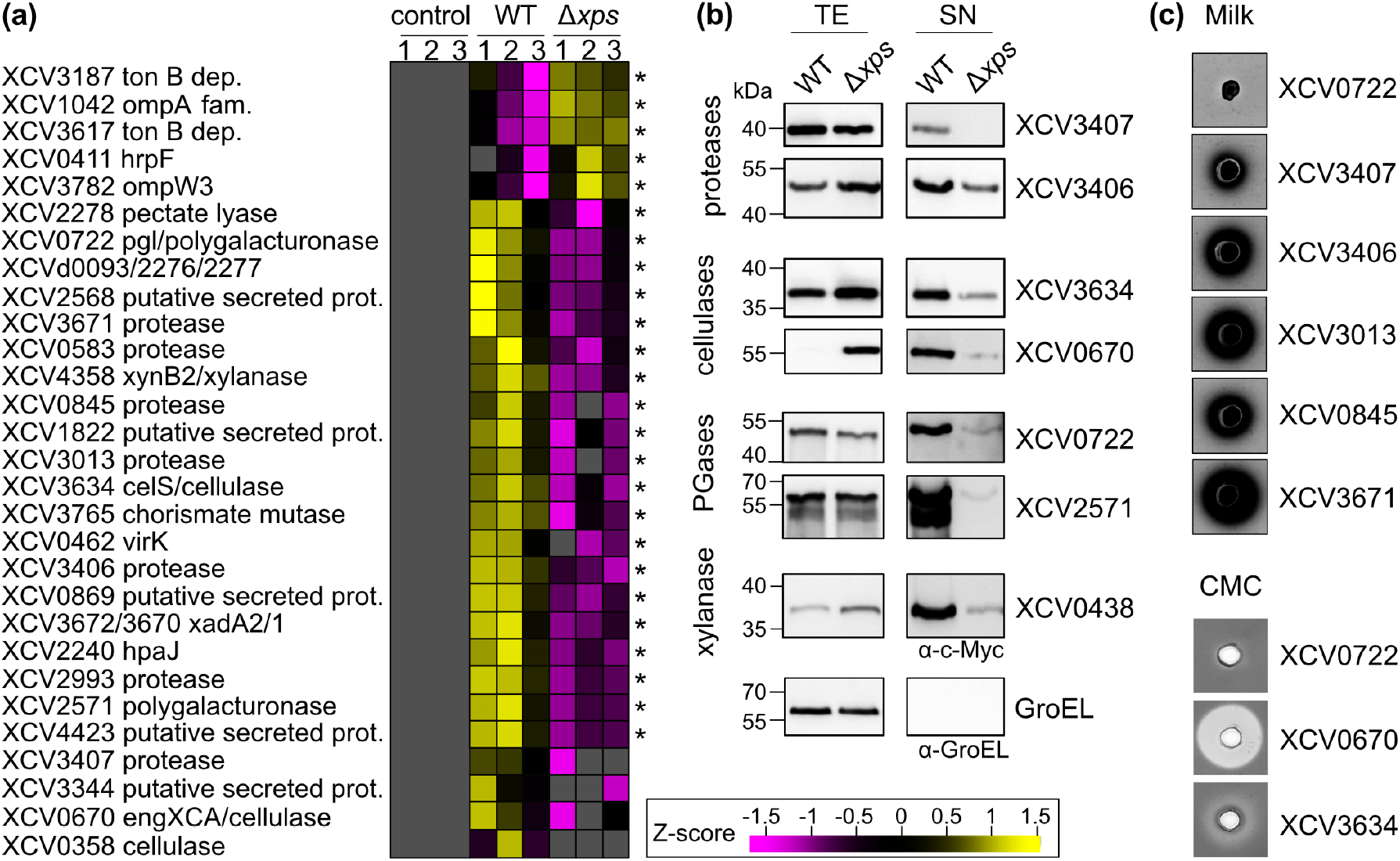
Differential quantitative *in planta* proteomics identified 22 putative T2Es including proteases CWDEs. **(a)** Heat map of normalized relative peptide abundance (assigned gene numbers and protein annotations on the left) showing data for peptides with consistent differential patterns across three independently infected plants (labeled 1, 2, 3). T2E candidates are those with higher Z-scores (yellow) during infection with wild-type (WT) *Xanthomonas* compared to the Δ*xps* mutant. Gray indicates no peptides were detected corresponding to a given protein. Asterisks denote statistically significant differences between conditions (Student’s t-test, p < 0.05). **(b)** *In vitro* secretion assays with candidate proteins in *Xe* strains grown in minimal media. Detection of c-Myc epitope-tagged proteins via SDS-PAGE and immunoblotting with a c-Myc epitope-specific antibody was performed on total extracts to confirm protein expression, while detection in the supernatant of cultures is indicative of their secretion. The cytoplasmic chaperonin GroEL was analysed as a lysis control. **(c)** Analysis of selected protease and cellulase activity of proteins expressed from *E.coli* in plate assays. Genes from *Xe* were overexpressed under control of a *lac* promoter in *E. coli* BL21 Star™ (DE3) cells. Strains were cultured in wells punched in LB plates containing 2% skim milk powder for protease activity or 1% carboxymethylcellulose (CMC) to assay for cellulase activity. Bacteria was incubated for 2 days at 28°C. The presence of a transparent halo surrounding the well is indicative of protease or cellulase activity. Halos on CMC plates were visualized via staining of CMC with 0.2% Congo Red. The putative polygalacturonase XCV0722 was used as a negative control in both experiments.

Overall, this strategy led to the identification of 20 novel *Xe* T2E candidate proteins (Fig. 7a). In total we identified 10 proteins annotated as carbohydrate-active enzymes (CAZymes; enzymes that assemble, modify, or break down carbohydrates; Drula *et al*. (2022)) targeting cellulose (three cellulases), hemicellulose (one xylanase), or pectin (one galactanase, two polygalacturonases, two pectate lyases). In addition to hydrolytic CAZymes, one carbohydrate binding protein of unknown function was detected. The identification of multiple pectin-degrading enzymes *in planta*, agrees with our observation that galacturonic acid is depleted from cell walls. Also fitting with the previously reported T2S-dependent protease activity of *Xe* on milk plates, we identified eight putative proteases: four serine proteases, one cysteine protease, and one metalloprotease, classified according to their catalytic mechanisms (MEROPS database, Rawlings *et al*. (2018)). Structural analysis using Foldseek (van Kempen *et al*., 2024) revealed XCV3407 similarity to fungal glutamic proteases. Additional T2E candidates included a putative alkaline phosphatase, an unknown protein, one chorismate mutase and the lytic murein transglycosylase HpaJ, which was previously shown to be coregulated with the T3S system (Noël *et al*., 2003).

### Validation of T2E candidates by *in vitro* secretion assays

To validate the proteomics approach, candidate T2Es were expressed as C-terminal c-Myc epitope-tagged derivatives under control of their native promoters in plant-mimicking XVM2 medium. As a positive control, the previously published T2E xylanase XCV4358 (Solé *et al*., 2015) was included, as well as the negative control, cytosolic GroEL. As expected, the levels of XCV4358 were reduced in Δ*xps* supernatants compared with those of the wild-type strain, confirming T2S-dependency. In contrast, GroEL was not detected in supernatants of wild-type or Δ*xps* strains, confirming minimal cell lysis. Like XCV4358, the identified proteases XCV3406 and XCV3407, the cellulases XCV3634 and XCV0670, and the polygalacturonases XCV0722 and XCV2571 were all detected in wild-type supernatants but substantially reduced in Δ*xps* supernatants (Fig. 7b), confirming T2S. Furthermore, XCV3406, XCV3634, XCV0670 and XCV4358 accumulated in cell extracts of the Δ*xps* mutant compared with those of the wild-type strain. The residual amounts of proteins, which were detected in culture supernatants of the Δ*xps* mutants, might be due to an alternative transport route for T2Es. Since T2Es have previously been detected in OMVs by immunoelectron microscopy (Solé *et al*., 2015), OMV-mediated release offers a plausible explanation for this residual secretion. The apparent discrepancy with prior T2S-independent secretion of XCV0670 and XCV0722 (Szczesny et al., 2010) is likely caused by different growth conditions; here we used native promoters and plant-mimicking medium which better reflects the *in planta* conditions. The T2S-dependent *in vitro* secretion of eight T2E candidates confirms that apoplastic proteomics is a suitable approach to identify *Xanthomonas* T2Es.

Enzymatic activity of a subset of T2Es was tested using milk and CMC plate assays in *E. coli* BL21 Star™ (DE3), which leaks periplasmic proteins into the medium (Zou *et al*., 2012). Cellulases XCV0670 and XCV3634 produced halos on CMC, confirming cellulase activity (XCV3634 halos were less pronounced). Proteases XCV3407, XCV3406, XCV3013, XCV3671 and XCV0845 all produced halos on milk plates (Fig. 7c). Polygalacturonase XCV0722 served as negative control.

### Genomic context of T2E candidates

To gain insight into the potential function and regulation of the identified T2E candidates, we examined whether any were encoded near genes that might be co-expressed as part of the same operon. Notably, the xylanase *xynB2* (XCV4358) gene lies within a predicted operon together with *XCV4357* (glucuronate metabolism, uxaC), *XCV4356* (a beta-galactosidase, bga2) and *XCV4355* (XynA). This operon is part of the *xytB* locus, which is conserved and characterized in *Xcc* (Dejean *et al*., 2013). The protease gene *XCV2993* shares a promoter with the gene encoding the ‘putative secreted protein’ (XCV2994). *XCV3671* is located 100 bp downstream of *XCV3672*, which encodes a putative secreted serralysin-like metalloprotease (UniProt, 2025). The gene encoding the non-peptidase serine protease homolog (XCV2568) is located 285 bp from the pectin lyase gene *XCV2569*. Other T2Es, showed no obvious evidence of operonic organization.

### Comparative analysis reveals differential conservation of T2Es

To place T2Es in a broader evolutionary context, we compared the complete predicted repertoire of periplasmic CAZymes across *Xcc*, *Xag* and *Xe*. CAZymes with secretion signals (SPCAZymes) were predicted across all three strains using SignalP 6.0 (Table S7). Of 69 SPCAZyme ortholog groups identified by OrthoFinder (Emms & Kelly, 2019), 41 (59%) were conserved across all three strains, consistent with a shared core enzymatic toolkit. However, 14 ortholog groups were shared exclusively between *Xag* and *Xe* but absent in *Xcc*, compared to only 3 shared exclusively between *Xcc* and *Xe*, suggesting greater overall similarity of the SPCAZyme repertoires. This is particularily notworthy in the context of a recent study identifying four *Xcc*-specific CWDEs; a β-glucanase (XC_0625), cellobiosidase CbsA (XC_0626), secreted lipase (XC_1740; not listed in the CAZyme database), and β-xylosidase (XC_3159), that together are required for *Xcc* to breach the hydathode-xylem tissue barrier and establish vascular infection (Paauw et al., 2024). Consistent with this, XC_0625, XC_0626 and XC_3159 appear in our SPCAZyme comparison as *Xcc*-conserved orthologs absent or divergent from those in the apoplast-colonizing *Xag* and *Xe*. Pectate lyase (PL) families showed particularly divergent distributions, with PL4 restricted to *Xag* and *Xe*, PL9 exclusive to *Xe*, and PL10 exclusive to *Xcc*). One confirmed T2E XCV2278 for example, belongs to the *Xe*-specific PL9 family. Although identification of putative orthologs across strains is suggestive of shared functions, expression *in planta* and secretion by the T2S system remains to be experimentally determined. Taken together, the broader SPCAZyme comparison supports the conclusion that core CAZyme families are conserved across these three *Xanthomonas* pathogens, but some lineage-restricted enzymes correlate with lifestyle (e.g. those identified by Paauw *et al*. (2024).

We also analyzed the identified T2E proteases in the context of all predicted periplasmic proteases (SP proteases) across all three strains (Table S8). Of the proteases identified as T2Es in this study, we observed distinct conservation patterns across the three strains. XCV2993 (S8 family) has two likely orthologs in *Xag* (XAR_1246, XAR_1244) and one in *Xcc* (XC_1450), although the latter has diverged more considerably. XCV0583 (S53 family) and XCV3671 (S8 famiily) have orthologs in *Xag* (XAR_1913) but no counterpart in *Xcc*. XCV3013 (cysteine protease, C family) has an ortholog in *Xcc* (XC_1422) but no counterpart in *Xag*. The last four, XCV0845 (S53 serine endopeptidase), XCV3406 (M72 metalloendopeptidase), XCV3407 (glutamic protease-like), and XCV2568 (non-peptidase serine protease homolog), all belong to *Xe*-specific ortholog groups with no counterpart in either *Xcc* or *Xag*. Taken together, there is greater similarity between *Xe* and *Xag* than between either and *Xcc*. However, the majority of *Xe* T2E proteases have no orthologous SP proteases in the other strains, suggesting the extracellular proteolytic activity is likely more strain specific.

## Discussion

A striking and initially unexpected outcome of this study is the demonstration that the T2S system contributes to *Xe* virulence in tomato to an extent comparable to that of the T3S system and that this contribution persists even in the absence of T3S activity. The additive phenotype of the double mutant confirms that both systems fulfill distinct and critical functions during infection. To define the nature of these T3S-independent functions, we investigated two non-mutually exclusive hypotheses: (i) T2Es contribute to virulence by suppressing plant immunity, and (ii) T2Es promote bacterial proliferation through nutrient acquisition from the plant cell wall. Although individual T2Es may have an undiscovered role in plant immunity, we found no evidence for their collective role in suppressing PTI when utilizing callose deposition as a readout. Rather, T2E activity may itself trigger immune responses that are normally suppressed by T3Es. This is consistent with findings in rice-infecting *Xanthomonas* where cell wall degradation products have been implicated in immune activation (Jha *et al*., 2007; Sinha *et al*., 2013). Based on these observations, we concluded that the primary T3E-independent contribution of the T2S system was likely related to nutrient acquisition.

### The apoplast as a nutritionally contested niche during infection

Carbon and nitrogen sources are present in the apoplast of healthy plants, however, the evolution of a plethora of bacterial and fungal pathogenic mechanisms that enhance apoplast sugar and amino acid content provides indirect evidence that nutrients are limiting during infection (Roussin-Léveillée *et al*., 2024). For example, in *Xanthomonas* species infecting rice, cotton, cassava, tomato and pepper, multiple independent T3Es converge on the transcriptional activation of SWEET family sucrose transporters, presumably increasing sucrose export to promote bacterial proliferation (Gupta *et al*., 2021; Roussin-Léveillée *et al*., 2024). Some xanthomonads also indirectly target the plant cell wall via T3Es: AvrHah1 from *X. gardneri* induces endogenous pectate lyase expression promoting apoplast hydration (Schwartz *et al*., 2017), while *X. citri* pv*. citri* PthA4 triggers ectopic expression of host CWDEs, releasing cell wall-derived sugars that feedback on bacterial gene expression to upregulate xylan metabolism pathways (Phan *et al*., 2025). Countering these strategies, plant immune responses are known to include the active retrieval of apoplastic sugars and restriction of amino acid export (Roussin-Léveillée *et al*., 2024), further limiting soluble nutrients in the apoplast. More recently, Wang *et al*. (2025) described an even more direct nutritional strategy in *Xanthomonas*: the T3E AvrBs2 acts as a synthetase inside host cells, utilizing host UDP-α-D-galactose to generate cyclic sugar phosphodiester termed ‘xanthosan’ that is exported to the apoplast and taken up by the bacteria as a dedicated carbon source.

The T2S system and secreted CWDEs are ancestral to the *Xanthomonas* lineage, being universally conserved across all sequenced species (Jacobs *et al*., 2015; Alvarez-Martinez *et al*., 2021), suggesting direct degradation of plant cell wall polymers as the original bacterial solution to this nutritional challenge, predating the more specialized T3E-mediated strategies that evolved later in some lineages. However, this remained to be tested experimentally. Although our limited strain sampling precludes broad generalizations, the ability of *Xe*, *Xag* and *Xcc* to employ the T2S system for the utilization of cell wall-derived carbon and milk protein under nutrient-limiting conditions supports this hypothesis.

### Evidence for strain-specific T2E mechanisms

The T2S systems of all strains appear to fulfill a role in nutrition, yet comparative analysis revealed significant differences in substrate utilization patterns. All strains grew well on xylan. This is consistent with previous reports showing that secreted xylanases contribute to plant infection and that xylan and xyloglucan utilization genes are conserved in *Xanthomonas* species (Jha *et al*., 2007; Dejean *et al*., 2013; Solé *et al*., 2015; Vieira *et al*., 2021; Phan *et al*., 2025). However, T2S system-dependence for xylan utilization varied significantly between strains. In culture *Xe* and *Xag* relied on the T2S system for xylan metabolism, while *Xcc* did not. This is in line with our previous demonstration that key *Xcc* xylanases, XCC4115 (homolog of *Xe* xylanase XCV4358) and XCC4118 (XCV0965 homolog), are secreted via alternative pathways (Solé et al., 2015), and points to functional redundancy in extracellular enzyme secretion. This divergence is particularly striking given that xylan is the preferred substrate across all three strains, meaning that while the nutritional target is conserved, the mechanisms employed to access it have diversified.

Strain-specific differences are also evident in cell wall digestion patterns. T2E activity during *Xe* infections caused galacturonic acid depletion, likely reflecting homogalacturonan degradation, accompanied by relative increases in rhamnose and galactose. In contrast, *Xcc* growth on pepper cell wall extracts *in vitro* released rhamnose and galactose in addition to oligogalacturonides (Vorholter *et al*., 2012), suggesting divergent pectin degradation pathways between species. Differences in pectinolytic enzyme repertoires may explain strain specific degradation patterns. This hypothesis is supported by our SPCAZyme ortholog comparison which revealed divergent pectate lyase (PL) family distributions between *Xe* and *Xcc* (Table S7). Differences may also reflect the net outcome of T2S-dependent degradation and host-specific differences in cell wall remodeling *in vivo* (Munzert & Engelsdorf, 2025).

Despite the observed T2S system-dependent reduction in galacturonic acid and the secretion of multiple pectin-degrading enzymes *in planta* by the *Xe* strain, none of the strains grew well on commercial pectin as a sole carbon source. Further, the growth observed was independent of the T2S system. This apparent contradiction raises the question of why bacteria invest in pectin degradation at all? One possible explanation is that the citrus peel pectin used in growth assays is not a suitable substrate. Citrus pectin is typically highly methylated (degree of methylation >50%; Singhal and Hulle (2022)), which limits accessibility to polygalacturonases and pectate lyases without prior de-esterification (Wormit & Usadel, 2018). More parsimoniously, *Xanthomonas* may degrade pectin not for direct metabolic use but to expose hemicellulose for subsequent metabolism. This is supported by transcriptomic analysis of *X. citri* during plant infection, which showed no induction of bacterial pectin metabolism genes despite active pectin degradation, but instead specific upregulation of xylan metabolism pathways (Phan et al., 2025).

Differences in substrate preferences suggest that while core nutritional functions are likely conserved, T2E repertoires have diversified substantially among strains and these differences likely extend beyond nutrition. A striking example is *Xcc*, a vascular pathogen whose T2E repertoire has been adapted to breach the hydathode-to-vasculature tissue barrier (Paauw *et al*., 2024). Of the four CWDEs responsible for this function, none were identified in our proteomics data, and *Xe* encodes only a single ortholog (LipA, 50% identity), suggesting that T2E specialization reflects infection lifestyle as much as host adaptation. Characterizing T2E repertoires at the strain level is therefore essential for understanding genomic differences.

### Proteomic identification reveals diversity of extracellular arsenals

To characterize the molecular basis of T2S system functionality in *Xe*, quantitative-comparative proteomics identified 22 T2E candidates targeting all major cell wall polysaccharide classes alongside a substantial protease complement, revealing a more extensive and functionally diversified secreted arsenal than previously appreciated. Several lines of evidence support the validity of this dataset: 1) recovery of known T2Es XCV4358 and XCV3671 (Solé *et al*., 2015), 2) the presence of periplasmic targeting signals in all identified proteins and 3) the identification of VirK and cysteine protease homologs previously found in rice apoplast infected with *Xanthomonas oryzae pv. oryzae* (Wang *et al*., 2013; Wang *et al*., 2017).

The identification of eight protease candidates was unexpected. Discovery of XCV3407, with structural similarity to fungal glutamic proteases, highlights that novel enzymatic classes remain to be discovered in well-studied pathogens. We confirmed enzymatic activity of five proteases *in vitro*, and T2S-dependent growth of *Xe* on milk protein provides functional evidence that protease secretion contributes to nitrogen acquisition. Together with cell wall polysaccharide utilization findings, we propose that the T2S system is a comprehensive nutritional system capable of exploiting both polysaccharide and protein components of the apoplast.

Near-complete conservation of the identified *Xe* T2Es in *Xag* (19/22) contrasts with more limited conservation in *Xcc* (12/22), consistent with phylogenetic distance. However, CWDE homologs were disproportionately well conserved in *Xcc* relative to overall repertoire conservation, reflecting selective pressure for enzymes fulfilling core nutritional functions. Notably, two pectate lyases were absent in *Xcc*, supporting strain-specific differences in pectin degradation. Protease repertoires showed greater divergence (3/8 *Xe* proteases in *Xcc*), potentially reflecting adaptation to distinct host environments or uncharacterised roles in pathogenesis.

### T2S/T3S synergy in apoplast remodeling

The T2S system and CWDEs predate the T3S system in the *Xanthomonas* lineage (Jacobs *et al*., 2015), and our data supports the hypothesis that they fulfill functions independently of T3S system activity. Nevertheless, since the subsequent acquisition of the T3S system and T3Es during the evolution of many pathogens, remodeling of the plant apoplast into a hydrated, nutrient-rich environment is likely the result of a complex and concerted interplay between T2Es and T3Es. The co-regulation of some T2Es with T3S system/T3E genes by the transcriptional regulators HrpG and HrpX (Büttner & Bonas, 2010) suggests these systems may be functionally linked rather than acting independently, although this remains to be investigated during infection.

Evidence for T2Es enhancing T3S system functionality takes multiple forms. First, we previously proposed that cell wall digestion could remove a physical barrier to T3S system penetration, facilitating T3E translocation into host cells (Szczesny *et al*., 2010). This is based on the observation that ETI-induced cell death is stronger during infections with wild-type *Xe* compared to a strain with deletion of the essential T2S system ATPase *xpsE*. These investigations were strategically conducted in resistant pepper plants, where the ETI-induced hypersensitive response (HR) provided an indirect measurement of T3E translocation efficiency. Whether CWDEs are directly responsible for this effect, however, remains to be investigated. Second, cell wall degradation products may signal increases in T3S system and T3E bacterial gene expression. This hypothesis was supported by *in vitro* studies in *X. oryzae* pv. *oryzae* and *X. citri* pv. *citri* showing that mono- and oligosaccharides induce T3S system genes, their master transcriptional regulator HrpX, and galactose and xylose metabolism pathways (Ikawa *et al*., 2018; Vieira *et al*., 2021; Tsuge & Ikawa, 2023).

In the reverse direction, T3Es enhance T2E function through at least two mechanisms. Our callose deposition experiments indicate that T2E activity triggers PTI responses that are suppressed by T3E-mediated immunity, thereby protecting T2E function during infection. This suppression may also have evolutionary consequences: by reducing immune-mediated fitness costs associated with T2E activity, suppression of PTI by T3Es could create a selective space for T2E diversification, contributing to the repertoire divergence observed across strains (Pena *et al*., 2024). Additionally, Phan et al. (2025) demonstrated that T3E-driven activation of host CWDEs releases cell wall-derived sugars that feedback on bacterial gene expression, upregulating T2-secreted CWDEs and establishing a T3E–T2E feedforward loop that amplifies cell wall degradation. Together, these findings suggest that T2S and T3S system activities are dynamically and reciprocally tuned during infection in ways that extend well beyond simple transcriptional co-regulation.

### Conclusion

This study establishes the T2S system as a comprehensive nutritional system for *Xanthomonas*, providing the first direct *in planta* evidence that T2E-mediated cell wall remodeling occurs during disease. Our findings reveal that while this nutrition function appears conserved across phylogenetically distinct strains, T2E repertoires have diversified substantially, likely reflecting strain-specific adaptations to distinct host environments or lifestyles. The emerging picture is one of dynamic coordination between the T2S system and T3S system: two systems that fulfill unique functions but also engage in reciprocal enhancement to remodel the apoplast and sustain bacterial proliferation. Given that the T2S system is universally conserved across the *Xanthomonas* lineage and makes substantial contributions to pathogenicity, it represents an underexplored but broadly applicable target for disease control strategies. Systematic characterization of T2E repertoires and T2S/T3S coordination during infection remain exciting frontiers that this study begins to address.

## Supporting information

Tabls S8

Script S1

## Acknowledgments

We thank Vivian Linke for her optimization of apoplast washing, Juliane Siebert for optimizing secretion assays and Susanne Kirsten, Nga Pham, Carsten Proksch, Sylvia Krüger and Dominika Thieme for technical support. We thank our gardening staff (Bianca Rosinski and Philipp Becker and ZMBP greenhouse staff). We thank Leif Eldon Erickson for advice on figures and Martin Schattat for reading the manuscript. Thanks to Tina Romeis for hosting JLE as a Walter Benjamin fellow at the Leibniz Institute for Plant Biochemistry (IPB, Halle). This work was supported by the Centre for Plant Molecular Biology (ZMBP, Tübingen) and the IPB and by DFG funding: Walter Benjamin Fellowship (ER 1024/1-1) awarded to JLE and individual grants to DB (BU 2145/6-2) and TE (EN 1071/3-1). SG was funded by grants from the German Academic Scholarship Foundation and a Graduate Scholarship of the State of Saxony-Anhalt. Authors used Claude (Anthropic) for language editing and text condensation. Authors take full responsibility for the integrity and accuracy of the published work.

## Author contributions

SG and TS contributed equally to data acquisition and data analysis and writing of manuscript drafts.

KSM-E and TE performed cell wall monosaccharide analysis;

TE also contributed to manuscript review and writing.

LH performed callose measurements and growth curves.

AKS and IK performed secretion assays.

SM contributed to experimental design and executed the proteomics pipeline.

SK performed transmission electron microscopy and cell wall thickness measurements.

DB and JLE contributed to data acquisition and proteomics analysis, supervised the project and wrote the manuscript.

## Data availability

The complete list of *Xanthomonas* peptides identified from the tomato apoplast *in planta* is available in Table S7. Strains generated in this study will be made available upon request. Raw mass spectrometry data will be deposited in the PRIDE repository upon publication.

## SUPPLEMENTARY MATERIAL

### Supplemental Material and Methods

- Bacterial strains and growth conditions
- Growth curves in modified XVM2 medium
- Plant material and plant inoculations
- Analysis of cell wall thickness
- Cell wall composition analyis
- Apoplast wash fluid sample preparation
- Identification of homologous CAZymes and proteases across *Xanthomonas* strains

### Supplementary Tables

**Table S1.** Bacterial strains and plasmids used in this study.

**Table S2.** Amino acid sequence conservation of *Xe* T2S system components in *Xag* and *Xcc*.

**Table S3.** Cell wall monosaccharide amounts in Arabidopsis, soybean, tomato and pepper leaf material.

**Table S4.** Monosaccharide composition of *Xe* grown *in vitro*.

**Table S5.** Cell wall monosaccharide composition of pepper leaves after infection with *Xe*.

**Table S6.** Plant cell wall thickness in pepper leaves in relation to bacterial strain and time point after infection

**Table S7.** *Xe* proteins from CAZy database with predicted signal peptides (Signal P-6.0).

**Table S8.** *Xanthomonas* proteins identified via mass spectrometry of apoplast wash fluid samples from infected plants.

**Table S9.** Comparison of periplasmic protease orthologs across *Xcc*, *Xag*, and *Xe* with gene trees.

### Other Supplementary Files

**Supplementary Script S1.** Fiji Macro for processing and quantification of callose deposits.

