## Supplementary Materials and Methods for "Defining the *Xanthomonas euvesicatoria* type II-secreted effector arsenal reveals key roles in bacterial nutrition and diversity across species"

### Bacterial strains and growth conditions

*Escherichia coli* cells were cultivated in Lysogeny broth (LB; 1% Bacto Tryptone; 0.5% Bacto Yeast Extract; 0.5% NaCl; pH 7.5; Bertani, 1951) at 37°C. *Xanthomonas* were grown in liquid nutrient-yeast-glycerol (NYG; 0.1% (w/v) Bacto Yeast Extract; 0.5% (w/v) Bacto Peptone; 0.2% (w/v) glycerol; pH 7.2; Daniels *et al.* (1984)) or plant mimicking media (XVM2; 20 mM NaCl; 10 mM $(NH_4)_2SO_4$; 5 mM $MgSO_4$; 1 mM $CaCl_2$; 0.16 mM $KH_2PO_4$; 0.32 mM $K_2HPO_4$; 0.01 mM $FeSO_4$; 10 mM fructose; 10 mM sucrose; 0.03% casaminoacids; pH 6.7; Wengelnik *et al.* (1996)). Plasmids were introduced into *E. coli* and *Xanthomonas* spp. by electroporation. Only pOGG2 plasmids were introduced into *Xanthomonas* by triparental mating.

### Growth curves in modified XVM2 medium

Growth experiments were performed in modified XVM2 minimal medium (Vieira *et al.*, 2021) in which either casamino acids were substituted with 3 g/L skim milk powder (XVM2 milk), or fructose and sucrose were substituted with cell wall extracts (AIR; alcohol insoluble residue, see below for extraction protocol) from plants at 0.5 mg/mL or isolated cell wall components: carboxymethylcellulose (CMC; Sigma-Aldrich, Steinheim) at 4 mg/mL, or pectin from citrus peel (Sigma-Aldrich, Steinheim), xylan from corn cob (Carl-Roth, Karlsruhe) or glucose (Carl Roth, Karlsruhe) at a concentration of 0.4 mg/mL. Exact composition of modified XVM2 mediate can be seen below.

Bacteria were grown overnight in NYG medium, cells were harvested and resuspended modified XVM2 media, washed once, and adjusted to 2 × $10^7$ cfu/mL. Cultures were grown shaking at 200 rpm at 30°C with bacterial density quantified every 30 min or after 24 h. For carbohydrates bacterial densities were quantified with a spectrophotometer after 5 min to allow for non-soluble AIR particles to settle.

**Plant material and plant inoculations**

*Solanum lycopersicum* cultivars (Heinz, VF26 and Moneymaker), *Glycine max* (ROYKA) and *Capsicum annuum* (Early Cal Wonder) were grown under long-day conditions (16 h light / 8 h dark) at 26°C (60% relative humidity) during the day and 19°C (40% relative humidity) at night. *Arabidopsis thaliana* (Oy-0) was grown under short-day conditions (8 h light / 16 h dark) at a constant 22°C and 50% relative humidity.

Bacteria were suspended in 10 mM $MgCl_2$ and adjusted to the specified densities. *Xe* was then infiltrated into third or fourth leaves on 4-6-week-old tomato plants and *Xag* was infiltrated into 4-week-old soybean leaves via a needless syringe. *Xcc* was introduced into *A. thaliana* vasculature via clip inoculation (Paauw *et al.*, 2024). Clip inoculations were incubated at 100% humidity for 16 h in the dark and then returned to standard growth conditions. For *in planta* growth curves bacteria were adjusted to $5 \times 10^4$ cfu/mL (Bonas *et al.*, 1991) and inoculated into the leaves of four plants. Two 4 mm leaf discs were collected and pooled at each time point (same leaf was used across time points). Samples were ground, diluted in series and plated on NYG plates containing rifampicin and cycloheximide. Bacterial colonies were quantified after 2-3 days.

For dip-infections *Xanthomonas* overnight cultures were washed twice in 10 mM $MgCl_2$, adjusted to $6 \times 10^7$ cfu/mL, supplemented with 0.02% Silwet L-77, and used to dip-infect 4-5 week-old tomato plants (30 s).

**Analysis of cell wall thickness**

Cell wall thickness was quantified from transmission electron micrographs in ImageJ (version 1.54, NIH; Schneider *et al.* (2012)) by manually measuring directly beneath sites of bacterial infection and in bacteria-free regions of the same cell (control). Statistical analysis was performed using SPSS 27 (IBM). As data were not normally distributed (Shapiro-Wilk test, $p < 0.05$), statistical comparisons were performed using the non-parametric Mann-Whitney U test.

**Cell wall composition analysis**

For cell wall analysis of un-infected leaves, samples were collected from 4-week-old *S. lycopersicum*, *G. max*, *C. annuum* and *A. thaliana*. For analysis of cell walls during infection, 4-week-old pepper leaves were inoculated with *Xanthomonas* via needless syringe at a bacterial density of $2 \times 10^7$ cfu/mL or a 1mM $MgCl_2$ buffer control. Leaves were collected five days postinfiltration. For analysis of *Xe* cell wall composition, bacteria were cultured in 50mL of NYG overnight to an $OD_{600}$ of approx. 1.5, and bacterial pellets were collected via centrifugation.

Plant samples were ground in liquid nitrogen prior to extraction. All samples were extracted in 80% ethanol at 80°C for 10 min under shaking and subsequently centrifuged at 17, 000×g for 10 min at room temperature. The process of removing alcohol soluble saccharides was repeated 3 times and the pellet was washed in 1 mL acetone to obtain alcohol insoluble residue (AIR). For analysis of cellulose and cell wall matrix monosaccharides in non-infected or *Xe*-infected plant material, dried AIR was subjected to acid hydrolysis with sulfuric acid as described by Yeats *et al.* (2016). Hydrolysis of *Xe* cell walls was performed by autoclaving bacterial AIR in 4% (w/v) sulfuric acid at 121°C for 60 min, after validating that pretreatment with 72 % (w/w) sulfuric acid for 1 h did not alter the release of monosaccharides.

All samples were diluted 30-fold in ultrapure water and 50 µM ribose was added to each sample as an internal standard. Quantification by high-performance anion-exchange chromatography with pulsed amperometric detection (HPAEC-PAD) was performed on a biocompatible Knauer Azura HPLC system equipped with Thermo Fisher Dionex CarboPac PA20 BioLC guard and analytical columns (3 × 30 mm and 3 × 150 mm) and coupled to an Antec Decade Elite SenCell detector as described by Reckleben *et al.* (2025). Calibration curves (1-100 µM) were measured for all monosaccharides investigated. To calculate relative changes in plant cell wall monosaccharide composition after *Xe* infection, the relative amount (mol %) of fucose, rhamnose, arabinose, galactose, xylose and galacturonic acid was subtracted from the mean mol % value of $MgCl_2$ infiltrated controls.

**Apoplast wash fluid sample preparation**

Acetone precipitation of proteins was performed by adding 800 µL of acetone to 200 µL of APW and incubated for 4 h at -20°C. Samples were then centrifuged at 4°C for 1 min at 4000×g and the acetone was removed. Pellets were resuspended in 75 µL of 2× Laemmli, boiled for 10 min, loaded on an 10% SDS gel and run to remove non-protein contaminants (Gonzalez *et al.*, 2012). Gels were stained with ‘Der Blaue Jonas’ Coomassie (Biozol) and lanes of the gel were cut into two pieces per sample. Proteins were digested in-gel with trypsin and peptides were desalted (Majovsky *et al.*, 2014). Peptides were separated using liquid chromatography C18 reverse phase chemistry (EASY-Spray column, 50 cm, Thermo Fisher Scientific; 80-min gradient, 2% to 45% acetonitrile in 0.1% FA, 250 nL/min) and analysed on an Orbitrap Fusion Lumos mass spectrometer (Thermo Fisher Scientific; 2 kV, capillary temperature of 305°C) using standard data-dependent acquisition (TOP15, HCD fragmentation).

Proteins were identified using MaxQuant v2.0.1.0 (Cox & Mann, 2008) with standard settings for label free quantification (LFQ) against the proteomes for tomato (ITAG4.1) and *Xe* (UniProt ID 316273) amended with common contaminants. Carbamidomethylation of cysteine (C) was set as fixed modification and oxidation of methionine (M) was tolerated as a variable modification. Fast LFQ was enabled with match between runs for all samples, the two gel pieces per sample were treated as different fractions and combined into one result file under standard settings. Resulting protein groups tables were processed with Perseus (Tyanova *et al.*, 2016) to remove contaminants, reverse hits and proteins only identified by site. LFQ data was filtered for a minimum of three valid values and to exclude tomato proteins, and then log2 transformed and presented as Z-scores. Student's T-tests were used to compare wildtype- and mutant-inoculated samples ($p<0.05$).

**Identification of homologous CAZymes and proteases across *Xanthomonas* strains**

CAZymes from *Xe*, *Xag*, and *Xcc* were extracted from the CAZy database (Drula *et al.* (2022); cazy.org) and sequences were used for predicting periplasmic localization with SignalP 6.0 (Teufel *et al.* (2022); DTU Health Tech) were used as input for ortholog identification and gene tree inference, performed using OrthoFinder v2.5.5 (Emms & Kelly, 2019) via the Galaxy Europe platform (usegalaxy.eu). All-versus-all sequence similarity searches were conducted using DIAMOND, and orthogroups were inferred using the MCL algorithm with an inflation parameter of 1.5. Gene trees were constructed using Dendroblast, which derives phylogenetic distances directly from DIAMOND pairwise similarity scores.

Full proteomes were collected from the Uniprot website (UniProt (2025); https://www.uniprot.org/) and Signal P 6.0 predictions were used to identify all proteins likely to be localized to the periplasm. From this list all proteins annotated as proteases were extracted and used for ortholog identification in the same manner as used for the CAZymes.
