## Supplementary material for "Defining the *Xanthomonas euvesicatoria* type II-secreted effector arsenal reveals key roles in bacterial nutrition and diversity across species": Figure S1

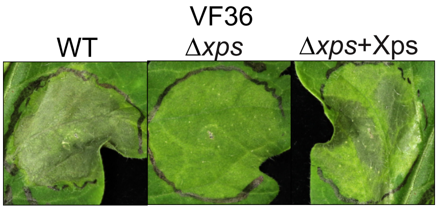


**Supplemental Fig. 1 The *Xe* type II secretion mutant shows reduced symptom formation in the VF36 cultivar.** VF36 tomato leaves were syringe-inoculated with *Xe* wild type strain 85-10, the T2S system mutant Δ*xps*, and the complementations strain Δ*xps*+Xps at an $OD_{600}$ of 0.05. Phenotypes of inoculations were documented at 6 dpi.
