## Supplementary material for "Defining the *Xanthomonas euvesicatoria* type II-secreted effector arsenal reveals key roles in bacterial nutrition and diversity across species": Tables S1-S6

**Table S1 Bacterial strains and plasmids used in this study.**

| Strains | Relevant characteristics[1] | References |
|---|---|---|
| ***Xanthomonas*** | | |
| *Xe* 85-10 | Pepper race 2, wild type; $Rif^R$ | Canteros (1990); Kousik and Ritchie (1998) |
| *Xe* 85-10Δ*hrcN* | In-frame deletion of the ATPase-encoding gene *hrcN*, $Rif^R$ | Lorenz and Büttner (2009) |
| *Xe* 85-10Δ*xps* | Derivative of strain 85-10 with a deletion of the entire *xps* gene cluster (nucleotide positions 4212581-4224331); $Rif^R$ | Goll et al. (2025) |
| *Xe* 85-10Δ*xps*Δ*hrcN* | Derivative of strain 85-10 lacking *hrcN* and the *xps* gene cluster; $Rif^R$ | This study |
| *Xe* 85-10Δ*xps* +Xps | *xps* deletion mutant complemented with the modulatar *xps* gene cluster on a plasmid; $Rif^R$ $Sm^R$ | Goll et al. (2025) |
| *Xe* 85-10Δ*xps*Δ*hrcN* +Xps | Derivative of strain 85-10 deleted in the *xps* gene cluster and *hrcN,* containing the modular *xps*-T2S gene cluster on a plasmid; $Rif^R$ | This study |
| *Xag* 8ra | Wild-type strain; $Rif^R$ | Hwang (1992) |
| *Xag* 8ra Δ*xps* | Derivative of strain *Xag* 8ra deleted in the entire *xps* gene cluster (nucleotide positions 4384268-4395300); $Rif^R$ | This study |
| *Xag* 8ra Δ*xps* +Xps | Derivative of strain *Xag* 8ra Δ*xps* complemented with the modulatar *xps* gene cluster on a plasmid; $Rif^R$ $Sm^R$ | This study |
| *Xcc* 8004 Δ*avrAC* | | |
| *Xcc* 8004 Δ*avrAC*Δ*xps* | Derivative of *Xcc* 8004 Δ*avrAC* deleted in the entire *xps* gene cluster (nucleotide positions 4230359-4241531); $Rif^R$ | This study |
| *Xcc* 8004 Δ*avrAC*Δ*xps* +Xps | Derivative of strain *Xcc* 8004 Δ*avrAC*Δ*xps* complemented with the modular *xps* gene cluster on a plasmid; $Rif^R$ $Sm^R$ | This study |
| ***E. coli*** | | |
| OneShot®TOP10 | $F^-$ *mcrA* Δ(*mrr*-hsdRMS-mcrBC) Φ80*lacZ*ΔM15 Δ*lacX74 recA1 araD139* Δ(*araleu*) 7697 *galU galK rpsL endA1 nupG*; $Str^R$ | Invitrogen |
| DH5α λpir | $F^-$ *recA hsdR17*($r_k^-$,$m_k^+$) Φ80d*lacZ DM15 [λpir]* | Menard et al. (1993) |
| BL21 StarTM (DE3) | F-*ompT hsdSB* (*rB-, mB-) gal dcm rne131* (DE3) | Invitrogen |
| **Plasmids** | | |
| **Complementation** | | |

| | | |
|---|---|---|
| pAGM8031 | Level M vector for modular Golden Gate cloning, derived from pBIN19 and pUC19, *lacZα* fragment flanked by *Bpi*I sites; $Sm^R$ | Addgene #48037; Weber et al., 2011 |
| pT2S038 | Derivative of pAGM8031 containing xpsE - xpsD with native promoters; $Sm^R$ | Goll et al. (2025) |
| **Mutagenesis** | | |
| pOGG2 | Golden Gate-compatible derivative of suicide vector pOK1, *sacB sacQ mobRK2 oriR6K*; $Sm^R$ | Schulze et al., 2012 |

| | | |
|---|---|---|
| pOGG2Δxps (Xe) | Derivative of pOGG2 carrying flanking regions of the *Xe xps* gene cluster (upstream: nucleotide position 4224332-4225081; downstream: nucleotide position 4212580-4213329) for deletion of xps gene cluster (nucleotide position 4212581-4224331) | Goll et al. (2025) |
| pOGG2Δ$xps_{Xag}$ | Derivative of pOGG2 carrying flanking regions of the *Xag xps* gene cluster (upstream: nucleotide position 4395301-4396300; downstream: nucleotide position 4383268-4384267) for deletion of the *xps* gene cluster (nucleotide position 4384268-4395300) | This study |
| pOGG2Δ$xps_{Xcc}$ | Derivative of pOGG2 carrying flanking regions of the *Xcc xps* gene cluster (upstream: nucleotide position 4241532-4242284; downstream: nucleotide position 4229609-4230358) for deletion of the *xps* gene cluster (nucleotide position 4230359-4241531) | This study |
| pRK2013 | ColE1 replicon, TraRK+ Mob+; $Km^R$ | Figurski & Helinski (1979) |
| **Secretion assays** | | |
| pBRM-P | Derivative of pBRM containing a *lacZα* fragment flanked by *Bsa*I recognition sites and a 3 x c-Myc epitope-encoding sequence lacking the *lac* promoter upstream of the 5' *Bsa*I site, $Gm^R$ | Szczesny et al. (2010) |
| pB-P3671 | pBRM-P derivative encoding XCV3671-3×c-Myc under control of the native *XCV3672* promoter (321 bp upstream of the predicted ATG start codon); $Gm^R$ | This study |
| pB-P4358 | pBRM-P derivative encoding XCV4358-3×c-Myc under control of the native *XCV4361* promoter (500 bp upstream of the predicted ATG start codon), $Gm^R$ | This study |
| pB-P0722 | pBRM-P derivative encoding XCV0722-3×c-Myc under control of the native *XCV0722* promoter (359 bp upstream of the predicted ATG start codon), $Gm^R$ | This study |
| pB-P0670 | pBRM-P derivative encoding XCV0670-3×c-Myc under control of the native *XCV0670* promoter (205 bp upstream of the predicted ATG start codon), $Gm^R$ | This study |
| pB-P3634 | pBRM-P derivative encoding XCV3634-3×c-Myc under the native XCV3634 promotor | This study |

| | | |
|---|---|---|
| | (477 bp upstream of the predicted ATG start codon), $Gm^R$ | |
| pB-P3406 | pBRM-P derivative encoding XCV3406-3×c-Myc under the native *XCV3406* promoter (390 bp upstream of the predicted ATG start codon), $Gm^R$ | This study |
| pB-P3407 | pBRM-P derivative encoding XCV3407-3×c-Myc under control of the native *XCV3407* promoter (613 bp), $Gm^R$ | This study |
| **T2E overexpression** | | |
| pB-XCV3671 | pBRM derivative encoding XCV3671-3×c-Myc under control of the *lac* promoter, $Gm^R$ | Szczesny et al. (2010) |
| pB-XCV3013 | pBRM derivative encoding XCV3013-3×c-Myc under control of the *lac* promoter, $Gm^R$ | Szczesny et al. (2010) |
| pB-XCV3406 | pBRM derivative encoding XCV3406-3×c-Myc under control of the *lac* promoter, $Gm^R$ | This study |
| pB-XCV3407 | pBRM derivative encoding XCV3407-3×c-Myc under control of the *lac* promoter, $Gm^R$ | This study |
| pB-XCV0845 | pBRM derivative encoding XCV0845-3×c-Myc under control of the *lac* promoter, $Gm^R$ | Solé et al., 2015 |
| pB-XCV0722 | pBRM derivative encoding XCV0722-3×c-Myc under control of the *lac* promoter, $Gm^R$ | This study |

[1]GM, gentamycin, Km, kanamycin; Rif, rifampicin; Sm, spectinomycin

**Table S2. Amino acid sequence conservation of *Xe* T2S system components in *Xag* and *Xcc.***

| Protein | *Xe* | *Xag* | *Xcc* |
|---|---|---|---|
| XpsE | XCV3668 (575 aa) | BIY41_18070 (575/575 aa; 99.3%; 2858) | XC3573 (560/575 aa; 97.4%; 2798) |
| XpsF | XCV3667 (405 aa) | BIY41_18065 (404/405 aa; 99.8%; 1996) | XC3572 (388/405 aa; 95.8%; 1934) |
| XpsG | XCV3666 (163 aa) | BIY41_18060 (143/143 aa; 100%; 733) | XC3571 (136/143 aa; 95.1%; 700) |
| XpsH | XCV3665 (169 aa) | BIY41_18055 (167/169 aa; 98.8%; 860) | XC3570 (146/169 aa; 86.4%; 740) |
| XpsI | XCV3664 (137 aa) | BIY41_18050 (129/138 aa; 93.5%; 658) | XC3569 (112/137 aa; 81.8%; 573) |
| XpsJ | XCV3663 (211 aa) | BIY41_18045 (176/180 aa; 97.8%; 887) | XC3568 (176/209 aa; 84.2%; 906) |
| XpsK | XCV3662 (301 aa) | BIY41_18040 (279/283 aa; 98.6%; 1403) | XC3567 (250/283 aa; 88.3%; 1265) |
| XpsL | XCV3661 (373 aa) | BIY41_18035 (364/373 aa; 97.6%; 1892) | XC3566 (342/373 aa; 91.7%; 1782) |
| XpsM | XCV3660 (217 aa) | BIY41_18030 (215/217 aa; 99.1%; 1076) | XC3565 (193/213 aa; 90.6%; 967) |
| XpsC | XCV3659 (265 aa) | BIY41_18025 (257/265 aa; 97.0 %; 1316) | XC3564 (226/265 aa; 85.3%; 1120) |
| XpsD | XCV3658 (759 aa) | BIY41_18020 (695/755 aa; 92.1%; 3496) | XC3563 (708/756 aa; 93.7%; 3572) |

Expasy pair-wise SIM-Alignments: BLOSUM 62; Gap open penalty 12; Gap extension penalty 4. Locus identifier (protein length in amino acids; % amino acid identity to the *Xe* homolog, and alignment score)

**Table S3. Cell wall monosaccharide amounts in 4-week-old Arabidopsis, soybean, tomato and pepper leaf material.**

| **[µg/mg AIR]** | ***Arabidopsis thaliana*** | ***Glycine max*** | ***Solanum lycopersicum*** | ***Capsicum annum*** |
|---|---|---|---|---|
| Fucose | 2.32 ± 0.29 | 7.24 ± 0.31 | 0.29 ± 0.01 | 0.50 ± 0.01 |
| Rhamnose | 5.82 ± 0.63 | 6.92 ± 0.28 | 5.32 ± 0.06 | 9.20 ± 0.38 |
| Arabinose | 9.08 ± 1.00 | 23.41 ± 1.24 | 7.54 ± 0.10 | 11.38 ± 0.42 |
| Galactose | 14.13 ± 1.43 | 16.71 ± 0.85 | 19.12 ± 0.44 | 18.09 ± 1.23 |
| Xylose | 7.89 ± 0.81 | 18.96 ± 0.86 | 9.28 ± 0.22 | 10.93 ± 0.60 |
| Mannose | 1.83 ± 0.26 | 7.36 ± 0.17 | 3.75 ± 0.05 | 3.51 ± 0.11 |
| Galacturonic acid | 36.58 ± 3.29 | 38.10 ± 4.11 | 37.95 ± 2.56 | 48.85 ± 2.86 |
| Glucuronic acid | 1.97 ± 1.15 | 1.06 ± 0.92 | 1.58 ± 0.74 | 0.25 ± 0.10 |
| Cellulose | 40.13 ± 3.52 | 61.32 ± 14.31 | 74.01 ± 8.54 | 66.26 ± 1.97 |

The amounts of fucose, rhamnose, arabinose, galactose, xylose, mannose, galacturonic acid, glucuronic acid and cellulose are shown in µg/mg AIR. Values are means of 3 biological replicates ± standard devation.

**Table S4. Monosaccharide composition of *Xe* grown *in vitro*.**

| [µg/mg AIR] | *Xanthomonas euvesicatoria* |
|---|---|
| Rhamnose | 0.25 ± 0.05 |
| Glucose | 16.96 ± 2.36 |
| Mannose | 2.75 ± 0.10 |
| Glucosamine | 3.53 ± 0.07 |

The amounts of rhamnose, glucose, mannose and glucuronic acid in *Xe* grown in NYG liquid media (shown in µg/mg AIR). Values are means of 3 biological replicates ± standard deviation.

**Table S5. Cell wall monosaccharide composition of pepper leaves after infection with *Xe*.**

| **[mol %]** | **$MgCl_2$** | **WT** | ***Δxps*** | ***Δxps+Xps*** |
|---|---|---|---|---|
| Fucose | 0.51 ± 0.01 | 0.62 ± 0.02 | 0.56 ± 0.02 | 0.61 ± 0.03 |
| Rhamnose | 10.17 ± 0.22 | 13.34 ± 0.50 | 11.26 ± 0.44 | 12.88 ± 0.54 |
| Arabinose | 11.31 ± 0.67 | 12.84 ± 1.30 | 14.41 ± 1.72 | 16.38 ± 0.92 |
| Galactose | 14.56 ± 1.09 | 18.74 ± 1.25 | 15.86 ± 0.63 | 19.94 ± 0.68 |
| Xylose | 8.21 ± 0.17 | 7.36 ± 0.47 | 7.00 ± 0.48 | 7.41 ± 0.83 |
| Galacturonic acid | 55.23 ± 2.04 | 47.09 ± 2.57 | 50.92 ± 2.16 | 42.77 ± 0.85 |

Pepper leaves were inoculated with *Xe* wild-type (WT), the T2S system mutant (*Δxps)*, the complementation strains carrying the xps-T2S gene cluster from *Xe* on a plasmid (*Δ*xps+Xps), or a 1mM $MgCl_2$ buffer control. The relative contents of fucose, rhamnose, arabinose, galactose, xylose and galacturonic acid were quantified from leaf AIR samples five days after inoculation (show in mol %). Values are means of 5 biological replicates ± standard deviation.

**Table S6. Plant cell wall thickness in pepper leaves in relation to bacterial strain and time point after infection.**

| | Wild-type *Xe* bacterial region | | | Wild- type *Xe* control region | | | test statistics | | |
|---|---|---|---|---|---|---|---|---|---|
| | Mean wall thickness (nm) | SD | N | Mean wall thickness (nm) | SD | N | U | z | p |
| 0hpi | 161,78 | 69,63 | 13 | 152,46 | 30,1 | 13 | 86 | 0,077 | 0,96 |
| 8hpi | 161,97 | 33,46 | 18 | 167,23 | 43,56 | 18 | 171 | 0,285 | 0,791 |
| 1dpi | 161,58 | 33,24 | 27 | 174,06 | 45,07 | 27 | 423 | 1,012 | 0,312 |
| 2dpi | 186,79 | 55,89 | 22 | 201,67 | 54,53 | 22 | 283 | 0,962 | 0,336 |
| 5dpi | 325,75 | 132,44 | 8 | 380 | 215,28 | 8 | 35 | 0,315 | 0,798 |

| | Δ*xps* bacterial region | | | Δ*xps* control region | | | test statistics | | |
|---|---|---|---|---|---|---|---|---|---|
| | Mean wall thickness (nm) | SD | N | Mean wall thickness (nm) | SD | N | U | z | p |
| 0hpi | 166,54 | 53,53 | 12 | 159,96 | 38,45 | 12 | 74 | 0,115 | 0,932 |
| 8hpi | 169,57 | 43,42 | 24 | 183,02 | 63,58 | 24 | 308,5 | 0,423 | 0,673 |
| 1dpi | 151,24 | 34,44 | 27 | 150,34 | 35,95 | 27 | 347 | -0,303 | 0,762 |
| 2dpi | 198,21 | 49,96 | 27 | 191,44 | 44,24 | 27 | 347 | -0,303 | 0,762 |
| 5dpi | 340,91 | 213,14 | 23 | 313,53 | 195,47 | 21 | 229 | -0,294 | 0,769 |

Values are given for regions beneath the bacteria and for corresponding control regions. The table presents mean values, standard deviations, sample sizes (N) and test statistics (Mann-Whitney U-Test) for each condition.
