## Supplementary material for "Defining the *Xanthomonas euvesicatoria* type II-secreted effector arsenal reveals key roles in bacterial nutrition and diversity across species": Tabls S8

**Table S8. Comparison of CAZymes across *X. campestris* pv. *campestris* (*Xcc*), *X. axonopodis* pv. *glycines* (*Xag*), and *X. euvesicatoria* (*Xe*).**

CAZymes from *Xe*, *Xag*, and *Xcc* predicted to localize to the periplasm (SignalP 6.0, DTU Health Tech) were used as input for ortholog identification and gene tree inference, performed using OrthoFinder v2.5.5 (Emms & Kelly, 2019). Gene trees were constructed using Dendroblast. CAZyme family annotations were obtained from cazy.org. Conservation is indicated by: green, conserved in all three strains; yellow, conserved in two strains; red, strain-specific. Gene accession numbers are shown in brackets, with *Xcc* genes in blue, *Xag* genes in green, and *Xe* genes in purple. Proteins identified as important for xylem infection by Pauaw *et al.* (2024) are indicated in blue as 'xylem invasion'. Previously identified T2Es are indicated in dark orange (Solé *et al.*, 2015), newly identified T2Es from this study are shown in yellow, both marked with 'T2S'.

| Family / OG | Conservation | *Xcc* gene | *Xag* gene | *Xe* gene | Gene tree |
|---|---|---|---|---|---|
| **GH2**<br>OG10 ; β-mannosidase | All 3 strains | XC_1218 | <br>FPK90_18780 | XCV3210 (*manB*)<br>XCV3430 | 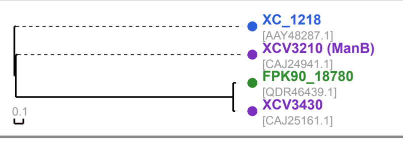 |
| **GH3**<br>OG1; β-glucosidase | All 3 strains | XC_1217<br>XC_4197<br>XC_2834<br>XC_3159 xylem invasion | FPK90_17225<br>FPK90_23575<br>FPK90_08085 | XCV3211<br>XCV4337<br>XCV1505 (*bglS*) | 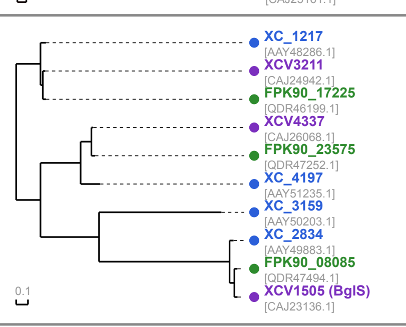 |
| **GH3**<br>OG16; 1,4-β-glucosidase | All 3 strains | XC_2461 [AAY49511.1] | FPK90_10155 [QDR45017.1] | XCV1823 (*celD*) [CAJ23500.1] | |
| **GH5**<br>OG3; cellulase | All 3 strains | XC_0026<br>XC_0027<br>XC_0028 | FPK90_00145<br>FPK90_00155<br>FPK90_00160 | XCV0029 (*egl1*)<br>XCV0031 (*egl2*)<br>XCV0033 (*egl3*) | 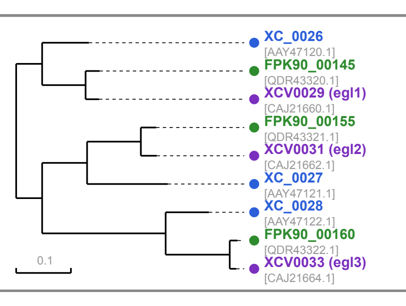 |
| **GH5**<br>OG17; endo-1,4-β-mannosidase | All 3 strains | XC_2458 [AAY49508.1] | FPK90_10175 [QDR45020.1] | XCV1826 [CAJ23503.1] | |
| **GH5**<br>OG49; cellulase | Xag + Xe | — | FPK90_02060 [QDR43635.1] | XCV0358 [CAJ21989.1] T2S | |
| **GH5,CBM2**<br>OG11; cellulase | All 3 strains | XC_0639<br>XC_0625 xylem invasion | FPK90_03550 | XCV0670 (*engXCA*) T2S | 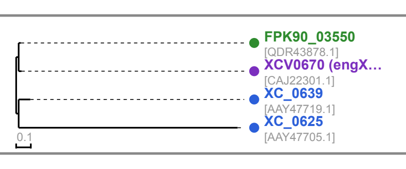 |
| **GH6**<br>1,4-β-cellobiosidase | Xcc only | XC_0626 [AAY47706.1] xylem invasion | — | — | |
| **GH8**<br>OG50; cellulase | Xag + Xe | — | FPK90_19820 (BcsZ) [QDR46621.1 | XCV3641 (*bcsZ*) [CAJ25372.1] | |
| **GH9**<br>OG18; cellulase | All 3 strains | XC_1727 [AAY48792.1] | FPK90_14415 [QDR45727.1] | XCV2704 (*egl4*) [CAJ24381.1] | |
| **GH10**<br>OG8; xylanase | All 3 strains | XC_4210 | FPK90_23685<br>FPK90_23695 | XCV4358 (*xynB2*) T2S<br>XCV4360 (*xynB3*) | 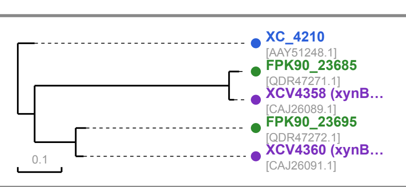 |
| **GH10**<br>OG19; endo-1,4-β-xylanase A | All 3 strains | XC_4207 [AAY51245.1] | FPK90_23665 [QDR47267.1] | XCV4355 (*xynA*) [CAJ26086.1] | |
| **GH12**<br>OG43; cellulase | Xcc + Xe | XC_0784 [AAY47862.1]<br>XC_0783 [AAY47861.1] | —<br>— | XCV3634 (*celS*) [CAJ25365.1] T2S<br>— | |
| **GH13**<br>OG51; α-amylase | Xag + Xcc | XC_3487 [AAY50530.1] | FPK90_04535 [QDR44046.1] | — | |
| **GH13**<br>OG63; cyclomaltodextrin glucanotransferase | Xcc + Xe | XC_1648 [AAY48714.1] | — | XCV2797 (*cgt*) [CAJ24476.1] | |
| **GH16**<br>OG53; endo-1,3-β-glucanase | Xag + Xcc | XC_3054 [AAY50102.1] | FPK90_07210 [QDR44505.1] | — | |
| **GH18**<br>OG27 | All 3 strains | XC_1220 [AAY48289.1] | FPK90_17210 [QDR46197.1] | XCV3208 [CAJ24939.1] | |
| **GH20**<br>β-hexosaminidase | Xcc only | XC_1219 [AAY48288.1] | — | — | |
| **GH23**<br>OG28; lytic murein transglycosylase | All 3 strains | XC_0758 [AAY47836.1] | FPK90_20080 [QDR46665.1] | XCV3686 (*slt*) [CAJ25417.1] | |
| **GH23**<br>OG29; lytic murein transglycosylase | All 3 strains | XC_1791 [AAY48854.1] | FPK90_14075 [QDR45666.1] | XCV2635 [CAJ24312.1] | |
| **GH23,CBM50**<br>OG30; membrane-bound murein hydrolase D | All 3 strains | XC_3259 [AAY50303.1] | FPK90_06165 [QDR44324.1] | XCV1086 (*mltD*) [CAJ22717.1] | |
| **GH23,CBM50**<br>OG54; lytic mureine transglycosylase | Xag + Xe | — | FPK90_15365 [QDR45880.1] | XCV2881 [CAJ24560.1] | |
| **GH28**<br>OG5; polygalacturonase | All 3 strains | XC_0705<br>XC_1849 | FPK90_03805<br>FPK90_13765 | XCV0722 (*pgl*) T2S<br>XCV2571 T2S | 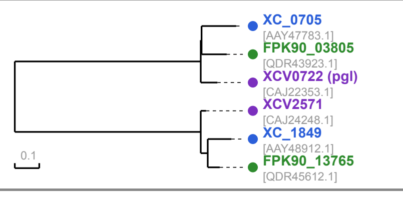 |
| **GH29**<br>OG55; α-L-fucosidase | Xag + Xcc | XC_2990 [AAY50038.1] | FPK90_07340 [QDR47491.1] | — | |
| **GH29**<br>OG31; α-L-fucosidase | All 3 strains | XC_1221 [AAY48290.1] | FPK90_17205 [QDR46196.1] | XCV3207 (*fucA1*) [CAJ24938.1] | |
| **GH30**<br>OG32; xylanase | All 3 strains | XC_3373 [AAY50417.1] | FPK90_06610 [QDR47477.1] | XCV0965 [CAJ22596.1] T2S | |
| **GH31**<br>OG33; α-xylosidase | All 3 strains | XC_2480 [AAY49530.1] | FPK90_10050 [QDR44998.1] | XCV1805 [CAJ23482.1] | |
| **GH33**<br>OG34; neuraminidase | All 3 strains | XC_1200 [AAY48269.1] | FPK90_17290 [QDR46209.1] | XCV3219 [CAJ24950.1] | |
| **GH35**<br>OG6; β-galactosidase | All 3 strains | XC_2481<br>XC_1708 | FPK90_10045<br>FPK90_14570 | XCV1804<br>XCV2736 | 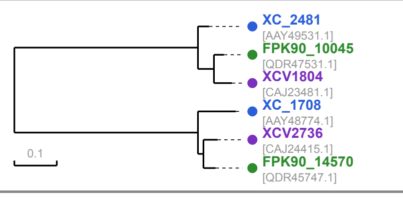 |
| **GH35**<br>OG35; β-galactosidase | Xag + Xcc only | XC_1214 [AAY48283.1]<br>— | FPK90_17265 [QDR46205.1]<br>FPK90_17235 [QDR46200.1] | —<br>— | |
| **GH37**<br>OG36; α-trehalase | All 3 strains | XC_0631 [AAY47711.1] | FPK90_03495 (TreA) [QDR43870.1] | XCV0660 (*treA*) [CAJ22291.1] | |
| **GH39**<br>OG56; β-xylosidase | Xag + Xcc | XC_4065 [AAY51104.1] | FPK90_22565 [QDR47084.1] | — | |
| **GH43**<br>OG57; xylosidase | Xag + Xe | — | FPK90_14515 [QDR45740.1] | XCV2728 [CAJ24407.1] | |
| **GH43,CBM91**<br>OG2; xylosidase | All 3 strains | XC_0158<br>XC_4154<br>XC_4196 | FPK90_00855<br>FPK90_23295<br>FPK90_23570<br>FPK90_07160 | XCV0150 (*xylB1*)<br>XCV4281 (*xylB2*)<br>XCV4336 (*xylB3*) | 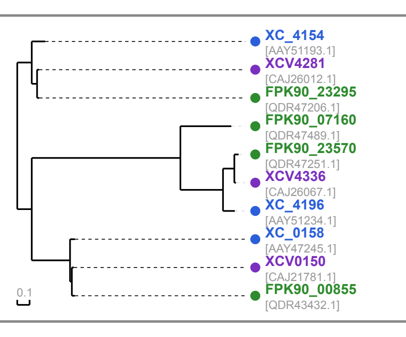 |
| **GH51**<br>OG37; α-L-arabinofuranosidase | All 3 strains | XC_3051 [AAY50099.1] | FPK90_07215 [QDR44506.1] | XCV1335 [CAJ22966.1] | |
| **GH53**<br>OG38; galactosidase | All 3 strains | XC_2984 [AAY50032.1] | FPK90_07355 [QDR44530.1] | XCV1360 [CAJ22991.1] | |
| **GH53**<br>OG39; arabinogalactan endo-1,4-β-galactosidase | All 3 strains | XC_0587 [AAY47668.1] | FPK90_03270 [QDR43840.1] | XCV0610 (*ganA*) [CAJ22241.1] | |
| **GH67**<br>OG58; α-glucuronidase | Xag + Xcc | XC_4193 [AAY51231.1] | FPK90_23555 [QDR47248.1] | | |
| **GH74**<br>OG40; cellulase | All 3 strains | XC_2483 [AAY49533.1] | FPK90_10035 [QDR44996.1] | XCV1802 [CAJ23479.1] | |
| **GH84**<br>OG41; β-N-acetylglucosaminidase | All 3 strains | XC_2155 [AAY49211.1] | FPK90_21990 [QDR46986.1] | XCV4043 [CAJ25774.1] | |
| **GH89**<br>No ortholog-α-N-acetylglucosaminidase | Xag only | — | FPK90_04050 [QDR43967.1] | — | |
| **GH92**<br>OG0; sugar hydrolase | All 3 strains | XC_3168<br>XC_1215<br>XC_4310<br>XC_0822 | FPK90_06630<br>FPK90_17260<br>FPK90_24190<br>FPK90_19550 | XCV1205<br>XCV3213<br>XCV4467 | 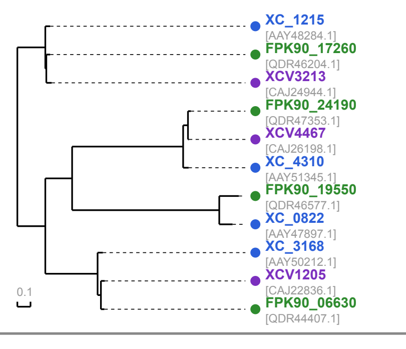 |
| **GH95**<br>OG13 | All 3 strains | XC_2479 | FPK90_10055<br>FPK90_13595 | <br>XCV2540 | 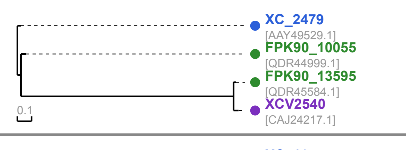 |
| **GH97**<br>OG7; α-glucosidase | All 3 strains | XC_1645<br>XC_1002 | FPK90_14880<br>FPK90_18785 | XCV2800<br>XCV3431 | 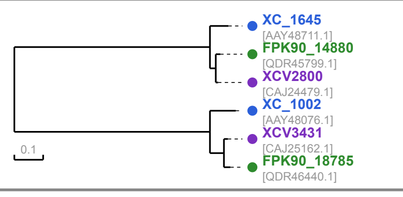 |
| **GH98,CBM35**<br>OG59; sugar binding | Xag + Xe | — | FPK90_04635 [QDR44064.1] | XCV0869 [CAJ22500.1] T2S | |
| **GH103**<br>OG9; transglycosylase | Xag + Xe | —<br>— | FPK90_23890<br>FPK90_15990<br>FPK90_18020 | XCV2440 (*hpaJ*) T2S<br>XCV4395 | 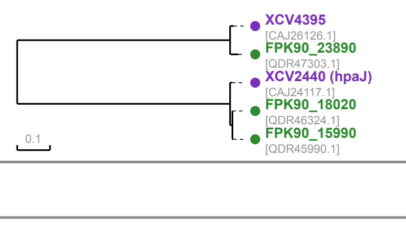 |
| **GH103**<br>OG20; membrane-bound lytic transglycosylase | All 3 strains | XC_0706 [AAY47784.1] | FPK90_03810 (MltB) [QDR43924.1] | XCV0723 (*mltB*) [CAJ22354.1] | |
| **GH113**<br>OG21 | All 3 strains | XC_3553 [AAY50596.1] | FPK90_19855 [QDR46626.1] | XCV3648 [CAJ25379.1] | |
| **GH115**<br>OG52 | Xag + Xe | — | FPK90_23365 [QDR47217.1] | XCV4294 [CAJ26025.1] | |
| **GH121**<br>OG22: carbohydrate binding | All 3 strains | XC_1713 [AAY48779.1] | FPK90_14520 [QDR47605.1] | BHE83_06305 [AOY69090.1] | |
| **GH125**<br>OG23 | All 3 strains | XC_1216 [AAY48285.1] | FPK90_17255 [QDR46203.1] | BHE83_08715 [AOY66637.1] | |
| **GH144**<br>OG24 | All 3 strains | XC_1911 [AAY48974.1] | FPK90_13440 [QDR45557.1] | XCV2509 [CAJ24186.1] | |
| **GH146**<br>OG25 | All 3 strains | XC_1718 [AAY48784.1] | FPK90_14500 [QDR45738.1] | BHE83_06285 [AOY66213.1] | |
| **GH158**<br>OG26 | All 3 strains | XC_3495 [AAY50538.1] | FPK90_04490 [QDR44039.1] | XCV0843 [CAJ22474.1] | |
| **GH186**<br>OG12; glucan biosynthesis | All 3 strains | XC_4237 | FPK90_23830 | BHE83_14405<br>XCV4387 (*opgD*) | 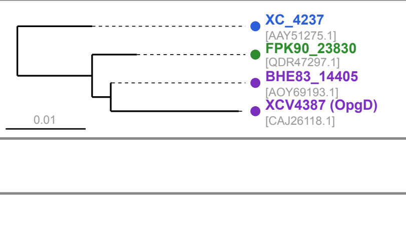 |
| **CE4,GH153**<br>No ortholog | Xag only | — | FPK90_10605 (PgaB) [QDR45088.1] | — | |
| **CE12**<br>OG15 | All 3 strains | XC_0163 [AAY47250.1] | FPK90_00895 [QDR43437.1] | XCV0155 [CAJ21786.1] | |
| **CE20**<br>OG4; 9-O-acetylesterase | All 3 strains | XC_2482<br>XC_4194 | FPK90_10040<br>FPK90_23560 | XCV1803<br>XCV4334 | 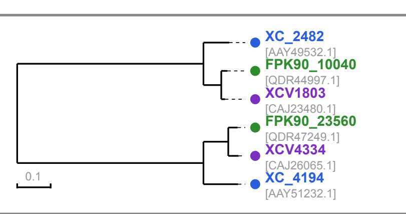 |
| **PL1**<br>OG14; pectate lyase | All 3 strains | AAY48367.1 | FPK90_16780<br>FPK90_13755 | <br>XCV3132 (*pel1*) | 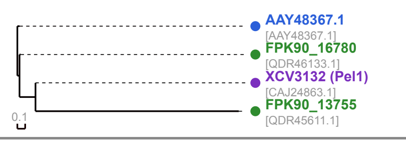 |
| **PL1**<br>OG44; pectate lyase | Xcc + Xe | XC_3591 [AAY50633.1]<br>XC_3590 [AAY50632.1] | —<br>— | XCV3687 (*pel2*) [CAJ25418.1]<br>— | |
| **PL4**<br>OG61; rhamnogalacturonase B | Xag + Xe | — | FPK90_19765 [QDR46612.1] | XCV3632 [CAJ25363.1] | |
| **PL9**<br>Pectate lyase | Xe only | — | — | XCV2278 [CAJ23955.1] T2S | |
| **PL10**<br>Pectate lyase | Xcc only | XC_0126 [AAY47217.1] | — | — | |
| **GHnc**<br>OG42; lytic transglycosylase | All 3 strains | XC_2042 [AAY49103.1] | FPK90_12505 [QDR45402.1] | XCV2379 [CAJ24056.1] | |
| **GHnc**<br>OG60; lytic transglycosylase | Xag + Xcc | XC_0707 [AAY47785.1] | FPK90_03815 [QDR43925.1] | — | |
| **CBM32**<br>OG45 | Xcc only | XC_1915 [AAY48978.1] | — | — | |
| **CBM50**<br>OG46; outer membrane lipoprotein | Xag + Xe | — | FPK90_09460 [QDR44899.1] | XCV1761 (*nlpD*) [CAJ23438.1] | |
| **CBM50**<br>OG47; peptidoglycan-binding | Xag + Xe | — | FPK90_21380 [QDR46883.1] | XCV3927 [CAJ25658.1] | |
| **Not present in the CAZyme database**<br>Arabinogalactan endo-beta-1,4-galactanase<br>Pectate lyase-like | | XC_1068 [AAY48138]<br>— | FPK90_18375 [QDR46383.1]<br>FPK90_24040 [QDR47328.1] | XCV3344 [CAJ25075.1] T2S<br>XCV4423 [CAJ26154.1] T2S | |


**Drula E, Garron M-L, Dogan S, Lombard V, Henrissat B, Terrapon N. 2022.** The carbohydrate-active enzyme database : functions and literature *Nucleic Acids Res* 50 : D571–D577.
**Emms DM and Kelly S. 2019.** OrthoFinder: phylogenetic orthology inference for comparative genomics. *Genome Biol* 20:238.
**Paauw M, Giesbers M, Pfeilmeier S, van den Burg HA. 2024.** Four cell wall-degrading enzymes of *Xanthomonas campestris* pv. *campestris* determine bacterial escape from hydathodes to the leaf vasculature. *BioRxiv preprint*.
**Solé M, Scheibner F, Hoffmeister AK, Hartmann N, Hause G, Rother A, Jordan M, Lautier M, Arlat M, Büttner D. 2015.** *Xanthomonas* pv. *vesicatoria* secretes proteases and xylanases via the Xps type II secretion system and outer membrane vesicles. *Journal of Bacteriology* 197(17): 2879–2893.
