## Supplementary material for "Defining the *Xanthomonas euvesicatoria* type II-secreted effector arsenal reveals key roles in bacterial nutrition and diversity across species": Table S9

**Table S9. Comparison of periplasmic protease orthologs across *X. campestris* pv. *campestris* (*Xcc*), *X. axonopodis* pv. *glycines* (*Xag*), and *X. euvesicatoria* (*Xe*) with gene trees.**

Protein sequences of proteases from *Xe*, *Xag*, and *Xcc* with signal peptides (SignalP 6.0; predicted periplasmic localization) were used as input for ortholog identification and gene tree inference, performed using OrthoFinder v2.5.5 (Emms & Kelly, 2019) via the Galaxy Europe platform (usegalaxy.eu). All-versus-all sequence similarity searches were conducted using DIAMOND, and orthogroups were inferred using the MCL algorithm with an inflation parameter of 1.5. Enzyme families are listed according to the MEROPS database (Rawlings *et al.*, 2018). Gene trees were constructed using Dendroblast, which derives phylogenetic distances directly from DIAMOND pairwise similarity scores. Protease family assignments were made using dbCAN. Conservation is indicated by: green, conserved in all three strains; yellow, conserved in two strains; red, strain-specific. Gene accession numbers are shown in brackets, with *Xcc* genes in blue, *Xag* genes in green, and *Xe* genes in purple. Previously identified T2Es are indicated in dark orange (Solé *et al.*, 2015), newly identified T2Es from this study are shown in yellow, both marked with 'T2S'.

<table>
<tr><th>Merops Family / OG</th><th>Conservation</th><th>Xcc gene</th><th>Xag gene</th><th>Xe gene</th><th>Gene tree</th></tr>
<tr><td colspan="6">Serine proteases</td></tr>
<tr><td>S1<br>OG13;DegP-like serine endoprotease</td><td>All 3 strains</td><td>XC_2972<br>[AAY50020.1]</td><td>XAR_0997<br>[EWC53554.1]</td><td>mucD<br>[CAJ23003.1]</td><td></td></tr>
<tr><td>S8<br>OG5;autrotransporter subtilase family</td><td>All 3 strains</td><td>XC_2159<br>XC_2942</td><td>XAR_0145<br>XAR_0967</td><td>XCV1402<br>XCV2109</td><td>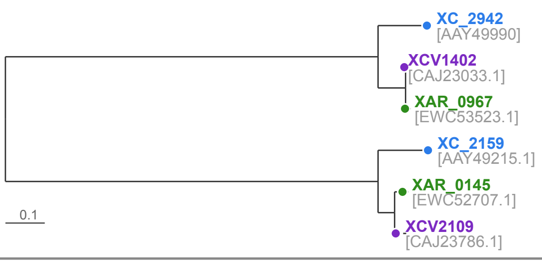
</td></tr>
<tr><td>S8<br>OG9; serine protease</td><td>All 3 strains</td><td>XC_1450</td><td>XAR_1246<br>XAR_1244</td><td>XCV2993 T2S</td><td>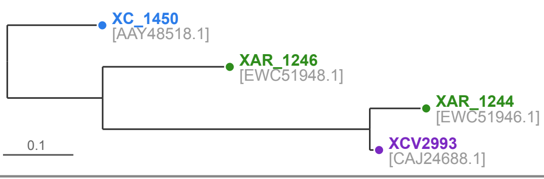
</td></tr>
<tr><td>S8<br>OG15; serine protease</td><td>All 3 strains</td><td>XC_1515<br>[AAY48583.1]</td><td>XAR_1325<br>[EWC52027.1]</td><td>XCV2918<br>[CAJ24597.1]</td><td></td></tr>
<tr><td>S8<br>OG1; serine protease</td><td>Xag + Xe</td><td></td><td>XAR_2311<br>XAR_2313<br>XAR_2314<br>XAR_0655</td><td>XCV0959<br>XCV0960<br>XCV0961<br>XCV3669<br>XCV3671 T2S</td><td>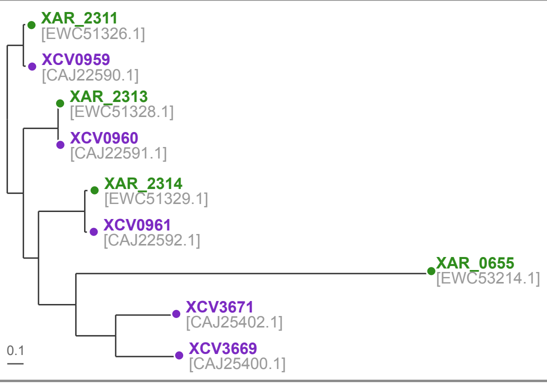
</td></tr>
<tr><td>S8<br>OG31; serine protease</td><td>All 3 strains</td><td>XC_3550<br>[AAY50593.1]</td><td>XAR_2933<br>[EWC50504.1]</td><td>XCV3639<br>[CAJ25370]</td><td></td></tr>
<tr><td>S9<br>OG0; dipeptidyl peptidase IV</td><td>All 3 strains</td><td>XC_4051<br>XC_1715<br>XC_1707</td><td>XAR_3955<br>XAR_1960<br>XAR_1530<br>XAR_1520</td><td>XCV4136<br>XCV2727<br>XCV2738</td><td>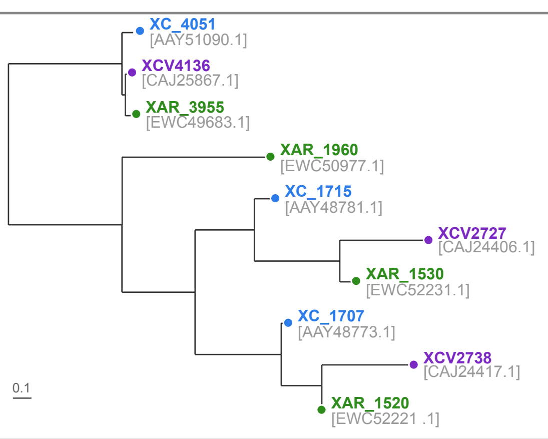
</td></tr>
<tr><td>S9<br>OG7</td><td>All 3 strains</td><td>XC_0253<br>XC_0844<br>XC_3481</td><td>XAR_3403</td><td>XCV0270<br>XCV1227</td><td>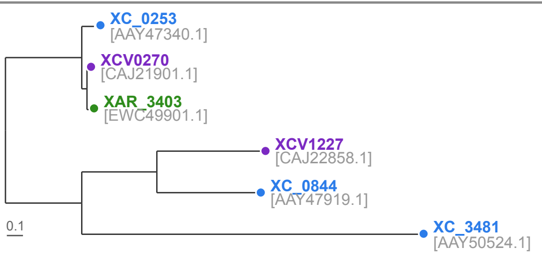
</td></tr>
<tr><td>S9<br>OG10</td><td>All 3 strains</td><td>XC_0654</td><td>XAR_1998<br>XAR_2001</td><td>XCV0685</td><td>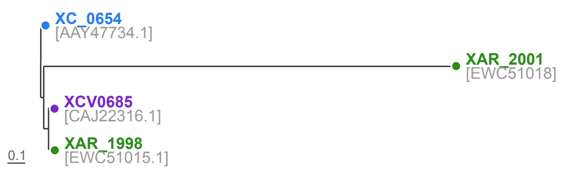
</td></tr>
<tr><td>S9<br>OG30</td><td>Xag + Xe</td><td></td><td>XAR_2599<br>[EWC51611.1]</td><td>XCV1232<br>[CAJ22863.1]</td><td></td></tr>
<tr><td>S10<br>OG11; serine carboxypeptidase</td><td>All 3 strains</td><td>XC_2538<br>[AAY49588.1]</td><td>XAR_0620<br>[EWC53179.1]</td><td>XCV1747<br>[CAJ23424.1]</td><td></td></tr>
<tr><td>S11<br>OG19; serine-type D-Ala-D-Ala carboxypeptidase</td><td>All 3 strains</td><td>XC_0708<br>[AAY47786.1]</td><td>XAR_2038<br>[EWC51054.1]</td><td>dacC<br>[CAJ22356.1]</td><td></td></tr>
<tr><td>S16<br>OG21</td><td>All 3 strains</td><td>XC_3271<br>[AAY50315.1]</td><td>XAR_2423<br>[EWC51437.1]</td><td>XCV1073<br>[CAJ22704.1]</td><td></td></tr>
<tr><td>S41<br>OG20</td><td>All 3 strains</td><td>XC_0714<br>[AAY47792.1]</td><td>XAR_2043<br>[EWC51059.1]</td><td>prc<br>[CAJ22361.1]</td><td></td></tr>
<tr><td>S41<br>OG33</td><td>Xag + Xe</td><td></td><td>XAR_3681<br>[EWC50178.1]</td><td>ctp<br>[CAJ21655.1]</td><td></td></tr>
<tr><td>S53<br>OG28; serine protease</td><td>Xag + Xe</td><td></td><td>XAR_1913<br>[EWC50930.1]</td><td>XCV0583 T2S<br>[CAJ22214.1]</td><td></td></tr>
<tr><td colspan="6">Metallo proteases</td></tr>
<tr><td>M1<br>OG18</td><td>All 3 strains</td><td>XC_0676<br>[AAY47755.1]</td><td>XAR_2016<br>[EWC51033.1]</td><td>pepN<br>[CAJ22331.1]</td><td></td></tr>
<tr><td>M1<br>OG6</td><td>All 3 strains</td><td>XC_3192<br>XC_0997<br>XC_0602</td><td>XAR_2552</td><td>pepN2<br>pepN3</td><td>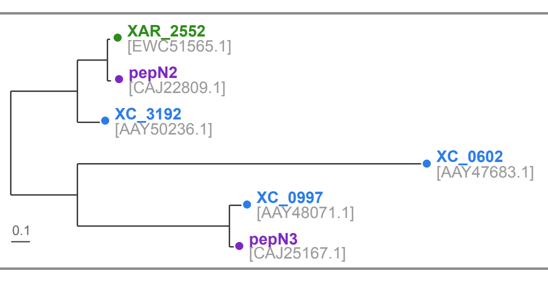
</td></tr>
<tr><td>M3<br>OG22</td><td>All 3 strains</td><td>XC_3130<br>[AAY50175.1]</td><td>XAR_2616<br>[EWC51628.1]</td><td>XCV1249<br>[CAJ22880.1]</td><td></td></tr>
<tr><td>M4<br>OG35</td><td>Xag + Xe</td><td></td><td>XAR_4207<br>[EWC49470.1]</td><td>XCV0495<br>[CAJ22126.1]</td><td></td></tr>
<tr><td>M4<br>OG36</td><td>Xcc + Xe</td><td>XC_0006<br>[AAY47100.1]</td><td></td><td>XCV0006<br>[CAJ21637.1]</td><td></td></tr>
<tr><td>M13<br>OG3; metalloendopeptidase</td><td>Xag + Xe</td><td></td><td>XAR_3138<br>XAR_1343<br>XAR_1341</td><td>XCV3834<br>XCV2898<br>XCV3027<br>XCV2900</td><td>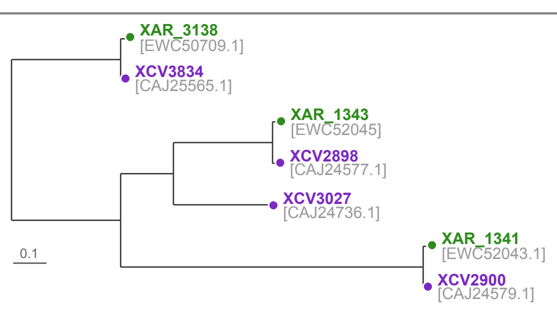
</td></tr>
<tr><td>M13<br>OG4</td><td>Xcc + Xag</td><td>XC_1406<br>XC_1542<br>XC_1543<br>XC_1544<br>XC_2148<br>XC_3742</td><td>XAR_1342</td><td></td><td>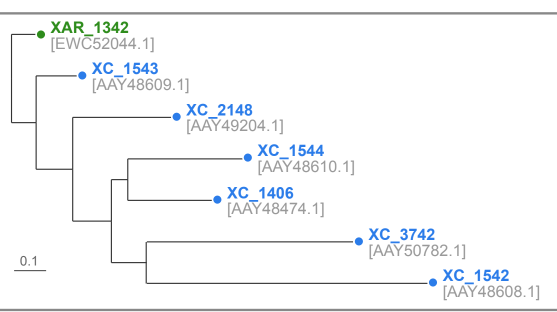
</td></tr>
<tr><td>M14<br>OG12</td><td>Xcc + Xag</td><td>XC_2833<br>[AAY49882.1]</td><td>XAR_0865<br>[EWC53422.1]<br>XAR_3184<br>[EWC50754.1]</td><td></td><td></td></tr>
<tr><td>M14<br>OG29</td><td>Xcc + Xag</td><td>XC_3280<br>[AAY50324.1]</td><td>XAR_2413<br>[EWC51427.1]</td><td></td><td></td></tr>
<tr><td>M16<br>OG17; zinc metalloprotease</td><td>All 3 strains</td><td>XC_0636<br>[AAY47716.1]</td><td>XAR_1982<br>[EWC50999.1]</td><td>XCV0666<br>[CAJ22297.1]</td><td></td></tr>
<tr><td>M19<br>OG24</td><td>All 3 strains</td><td>XC_4317<br>[AAY51352.1]</td><td>XAR_3717<br>[EWC50214.1]</td><td>XCV4474<br>[CAJ26205.1]</td><td></td></tr>
<tr><td>M20<br>OG2</td><td>All 3 strains</td><td>XC_0643<br>XC_0024</td><td>XAR_1988<br>XAR_4002<br>XAR_3035</td><td>XCV0673<br>XCV4096<br>XCV0027</td><td>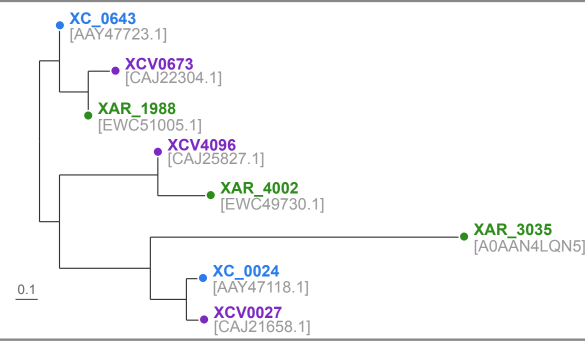
</td></tr>
<tr><td>M20<br>OG23</td><td>All 3 strains</td><td>XC_1008<br>[AAY48082.1]</td><td>XAR_2721<br>[EWC50292.1]</td><td>XCV3425<br>[CAJ25156.1]</td><td></td></tr>
<tr><td>M20<br>OG25</td><td>All 3 strains</td><td>XC_4105<br>[AAY51144.1]</td><td>XAR_3890<br>[EWC49620.1]</td><td>XCV4194<br>[CAJ25925.1]</td><td></td></tr>
<tr><td>M23<br>OG32</td><td>Xag + Xe</td><td></td><td>XAR_3680<br>[EWC50177.1]</td><td>XCV0025<br>[CAJ21656.1]</td><td></td></tr>
<tr><td>M28<br>OG8</td><td>Xag + Xe</td><td></td><td>XAR_3746<br>XAR_2663<br>XAR_3533</td><td>XCV4437<br>XCV1311</td><td>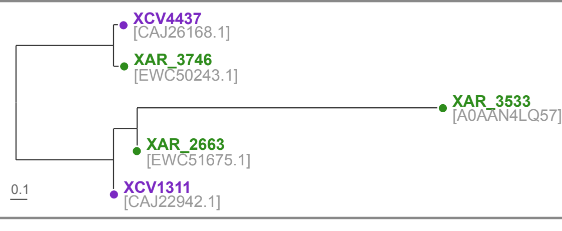
</td></tr>
<tr><td>M28<br>OG27</td><td>Xag + Xe</td><td></td><td>XAR_1083<br>[EWC51785.1]</td><td>XCV3148<br>[CAJ24879.1]</td><td></td></tr>
<tr><td>M23<br>OG14</td><td>Xag + Xe</td><td></td><td>XAR_1051<br>[EWC51753.1]</td><td>XCV3178<br>[CAJ24909.1]<br>XCVd0033<br>[CAJ19845.1]</td><td></td></tr>
<tr><td>M23<br>OG16</td><td>All 3 strains</td><td>XC_1857<br>[AAY48920.1]</td><td>XAR_1680<br>[EWC52380.1]</td><td>XCV2560<br>[CAJ24237.1]</td><td></td></tr>
<tr><td>Unassigned<br>OG37</td><td>Xcc + Xe</td><td>XC_1292<br>[AAY48361.1]</td><td></td><td>XCV3139<br>[CAJ24870.1]</td><td></td></tr>
<tr><td colspan="6">Cysteine proteases</td></tr>
<tr><td>Unassigned<br>OG34</td><td>Xcc + Xag</td><td>XC_4060<br>[AAY51099.1]</td><td>XAR_3945<br>[EWC49674.1]</td><td></td><td></td></tr>
<tr><td>C1<br>OG38</td><td>Xcc + Xe</td><td>XC_1422<br>[AAY48490.1]</td><td></td><td>XCV3013 T2S<br>[CAJ24716.1]</td><td></td></tr>
<tr><td colspan="6">Strain-specific (no ortholog)</td></tr>
<tr><td>S9</td><td>Xag only</td><td></td><td>XAR_1960<br>[EWC50977.1]<br>XAR_3415<br>[EWC49913.1]</td><td></td><td></td></tr>
<tr><td>Various</td><td>Xcc only</td><td>XC_1302<br>[AAY48371.1]<br>XC_2385<br>[AAY49438.1]<br>XC_3377<br>[AAY50422.1]<br>XC_3379<br>[AAY50423.1]<br>XC_3534<br>[AAY50577.1]<br>XC_3575<br>[AAY50617.1]<br>XC_3687<br>[AAY50728.1]<br>ddpX<br>[AAY50913.1]</td><td></td><td></td><td></td></tr>
<tr><td>Various</td><td>Xe only</td><td><br><br>S53; serine-type endopeptidase<br><br><br><br>similar to a glutamic protease<br>M72; metalloendopeptidase<br>non-peptidase serine protease homolog</td><td></td><td>XCV0161<br>[CAJ21792.1]<br>XCV0254<br>[CAJ21885.1]<br>XCV0845 T2S<br>[CAJ22476.1]<br>XCV1883<br>[CAJ23560.1]<br>XCV3964<br>[CAJ25695.1]<br>XCVd0044<br>[CAJ19856.1]<br>XCV3407 T2S<br>[CAJ25138.1]<br>XCV3406 T2S<br>[CAJ25137.1]<br>XCV2568 T2S<br>[CAJ24245.1]</td><td></td></tr>
</table>

**Emms DM and Kelly S. 2019.** OrthoFinder: phylogenetic orthology inference for comparative genomics. *Genome Biol* 20:238.

**Solé M, Scheibner F, Hoffmeister AK, Hartmann N, Hause G, Rother A, Jordan M, Lautier M, Arlat M, Büttner D. 2015.** *Xanthomonas* pv. *vesicatoria* secretes proteases and xylanases via the Xps type II secretion system and outer membrane vesicles. *Journal of Bacteriology* **197**(17): 2879–2893.

**Rawlings ND, Barrett AJ, Thomas PD, Huang X, Bateman A & Finn RD. 2018.** The MEROPS database of proteolytic enzymes, their substrates and inhibitors in 2017 and a comparison with peptidases in the PANTHER database. Nucleic Acids Res 46, D624-D632.
